# Bimodal inference in humans and mice

**DOI:** 10.1101/2021.08.20.457079

**Authors:** Veith Weilnhammer, Heiner Stuke, Kai Standvoss, Philipp Sterzer

## Abstract

2

Perception is known to cycle through periods of enhanced and reduced sensitivity to external information. Here, we asked whether such infra-slow fluctuations arise as a noise-related epiphenomenon of limited processing capacity or, alternatively, represent a structured mechanism of perceptual inference. Using two large-scale datasets, we found that humans and mice waver between alternating intervals of externally- and internally-oriented modes of sensory analysis. During external mode, perception aligned more closely with the external sensory information, whereas internal mode was characterized by enhanced biases toward perceptual history. Computational modeling indicated that dynamic changes in mode are enabled by two interlinked factors: (i), the integration of subsequent inputs over time and, (ii), infra-slow anti-phase oscillations in the perceptual impact of external sensory information versus internal predictions that are provided by perceptual history. Simulated data suggested that between-mode fluctuations may benefit perception by generating unambiguous error signals that enable robust learning and metacognition in volatile environments.

**One sentence summary:** Humans and mice fluctuate between external and internal modes of sensory processing.

## 4 Introduction

The capacity to respond to changes in the environment is a defining feature of life^1–3^. Intriguingly, the ability of living things to process their surroundings fluctuates considerably over time^4,5^. In humans and mice, perception^6–12^, cognition^13^ and memory^14^ cycle through prolonged periods of enhanced and reduced sensitivity to external information, suggesting that the brain detaches from the world in recurring intervals that last from milliseconds to seconds and even minutes^4,5^. Yet breaking from external information is risky, as swift responses to the environment are often crucial to survival.

What could be the reason for these fluctuations in perceptual performance^11^? First, periodic fluctuations in the ability to parse external information^11, 15, 16^ may arise simply due to bandwidth limitations and noise. Second, it may be advantageous to actively reduce the costs of neural processing by seeking sensory information only in recurring intervals^5,17^, otherwise relying on random or stereotypical responses to the external world. Third, spending time away from the ongoing stream of sensory inputs may also reflect a functional strategy that facilitates flexible behavior and learning^18^: Intermittently relying more strongly on information acquired from past experiences may enable agents to build up stable internal predictions about the environment despite an ongoing stream of external sensory signals^19^. By the same token, recurring intervals of enhanced sensitivity to external information may help to detect changes in both the state of the environment and the amount of noise that is inherent in sensory encoding^19^.

In this work, we sought to elucidate whether periodicities in the sensitivity to external information represent an epiphenomenon of limited processing capacity or, alternatively, result from a structured and adaptive mechanism of perceptual inference. To this end, we analyzed two large-scale datasets on perceptual decision-making in humans^20^ and mice^21^. When less sensitive to external stimulus information, humans and mice showed stronger serial dependencies^22–33^, which have been conceptualized as internal predictions that reflect the auto-correlation of natural environments^34^ and bias perceptual decisions toward preceding choices^30, 31, 35^. Computational modeling indicated that ongoing changes in perceptual performance may be driven by systematic fluctuations between externally- and internally-oriented modes of sensory analysis. Model simulations suggested that such bimodal inference may improve, (i), the ability to robustly determine the statistical properties of volatile environments and, (ii), the ability to calibrate internal beliefs about the degree of noise inherent in the encoding of sensory information.

## 5 Results

### 5.1 Human perception fluctuates between epochs of enhanced and reduced sensitivity to external information

We began by selecting 66 studies from the Confidence Database^20^ that investigated how human participants (N = 4317) perform binary perceptual decisions (Figure 1A; see Methods section for details on inclusion criteria). As a metric for perceptual performance (i.e., the sensitivity to external sensory information), we asked whether the participant’s response and the presented stimulus matched (*stimulus-congruent* choices) or differed from each other (*stimulus-incongruent* choices; Figure 1B and C) in a total of 21.05 million trials.

**Figure 1.**
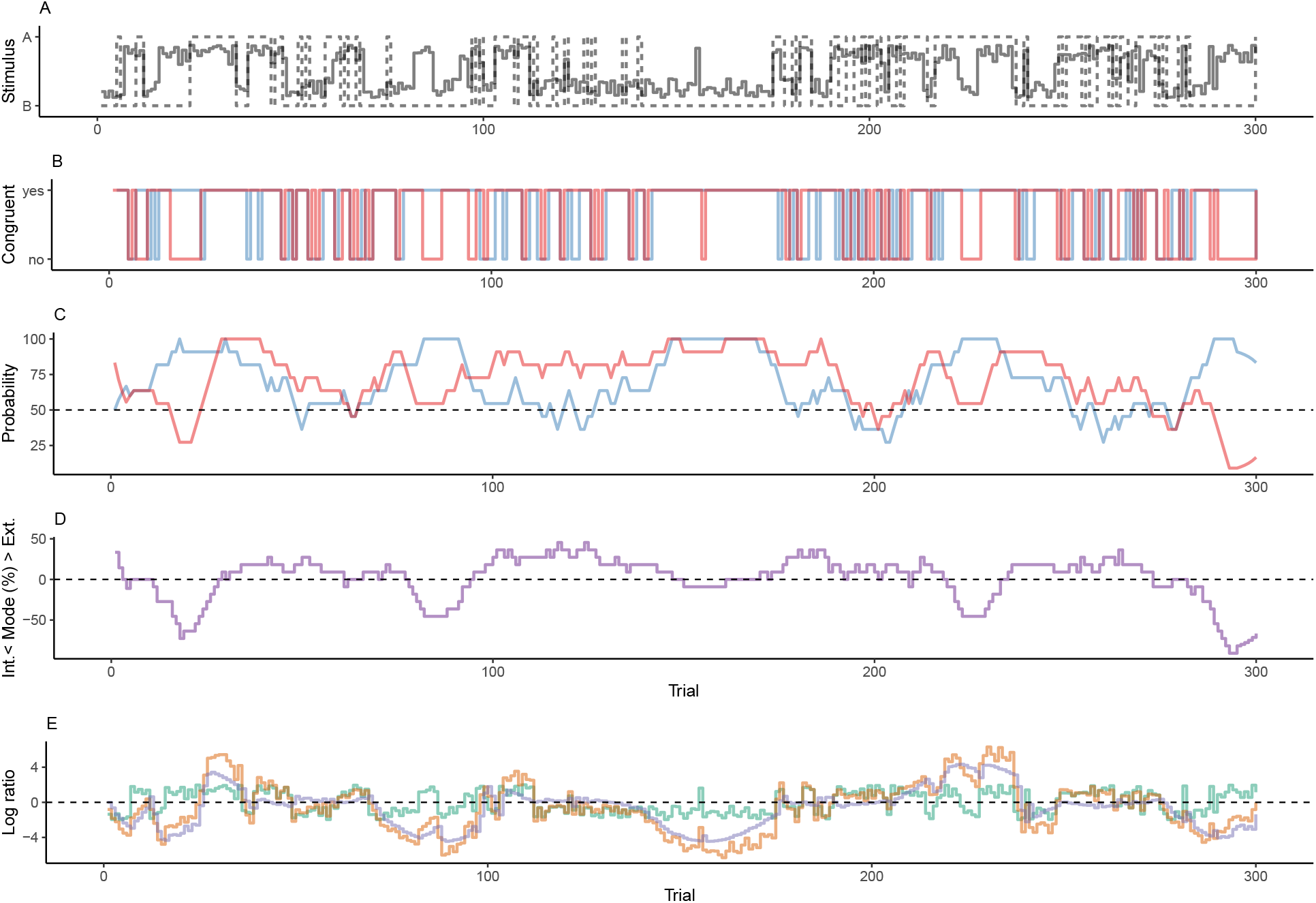
Concept. A. In binary perceptual decision-making, a participant is presented with stimuli from two categories (A vs. B; dotted line) and reports consecutive perceptual choices via button presses (sold line). All panels below refer to this example data. B. When the response matched the external stimulus information (i.e., overlap between dotted and solid line in panel A), perceptual choices are *stimulus-congruent* (red line). When the response matches the response at the preceding trial, perceptual choices are *history-congruent* (blue line). C. The dynamic probabilities of stimulus- and history-congruence (i.e., computed in sliding windows of ±5 trials) fluctuate over time. D. The *mode* of perceptual processing is derived by computing the difference between the dynamic probabilities of stimulus- and history-congruence. Values above 0% indicate a bias toward external information, whereas values below 0% indicate a bias toward internal information. E. In computational modeling, internal mode is caused by an enhanced impact of perceptual history. This causes the posterior (black line) to be close to the prior (blue line). Conversely, during external mode, the posterior is close to the sensory information (log likelihood ratio, red line).

In a first step, we asked whether the ability to accurately perceive sensory stimuli is constant over time or, alternatively, fluctuates in periods of enhanced and reduced sensitivity to external information. We found perception to be stimulus-congruent in 73.46% ± 0.15% of trials (mean ± standard error of the mean; Figure 2A), which was highly consistent across the selected studies (Supplemental Figure S1A). In line with previous work^8^, we found that the probability of stimulus-congruence was not independent across successive trials: At the group level, stimulus-congruent perceptual choices were significantly autocorrelated for up to 15 trials. Autocorrelation coefficients decayed exponentially over time (rate *γ* = −1.92 × 10^-3^ ± 4.5×10^-4^, T(6.88 ×10^4^) = −4.27, p = 1.98 × 10^-5^; Figure 2B). Importantly, the autocorrelation of stimulus-congruent perception was not a trivial consequence of the experimental design, but remained significant when controlling for the trial-wise autocorrelation of task difficulty (Supplemental Figure S2A) or the sequence of presented stimuli (Supplemental Figure S2B).

**Figure 2.**
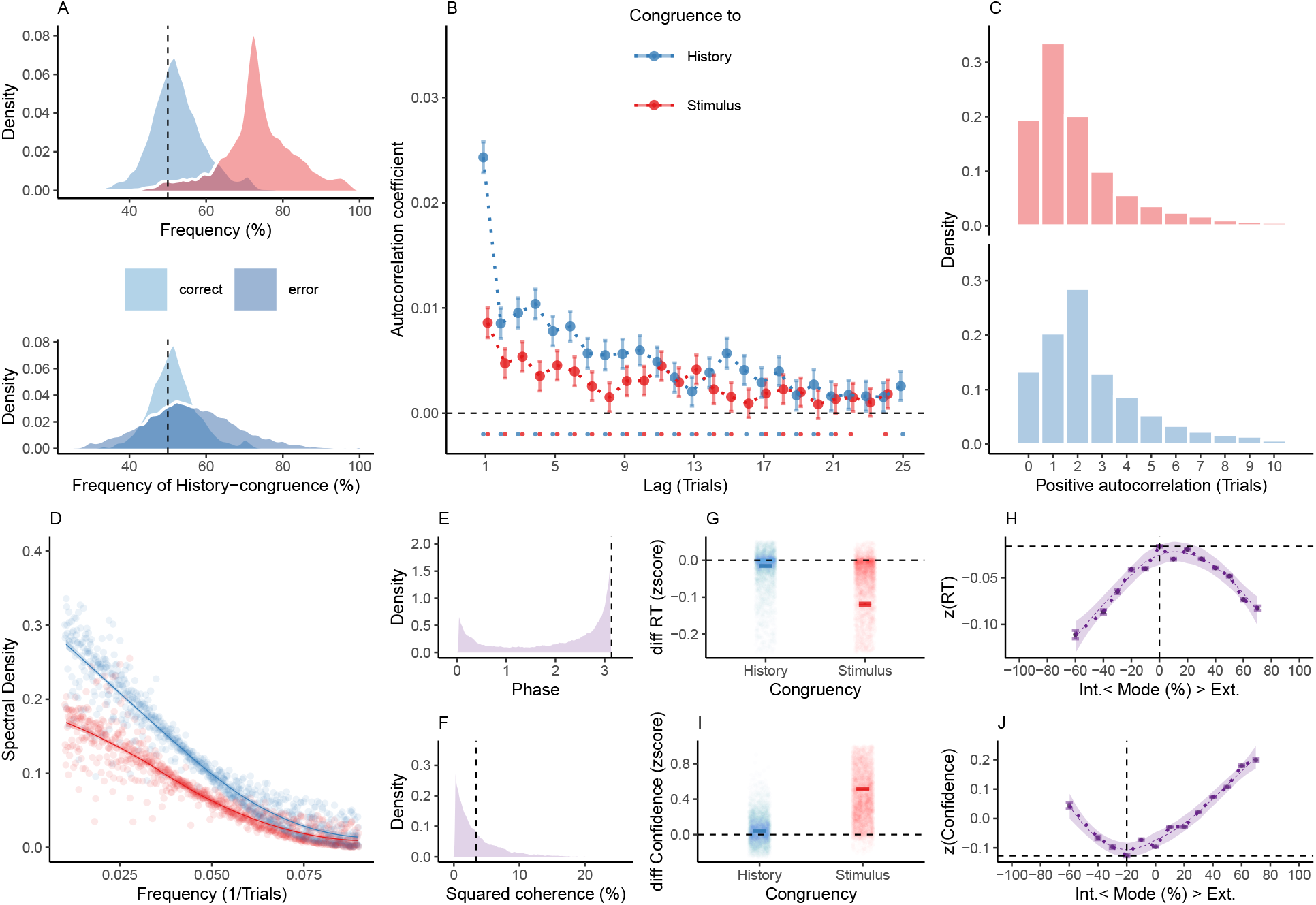
Internal and external modes in human perceptual decision-making. A. In humans, perception was stimulus-congruent in 73.46% ± 0.15% (in red) and history-congruent in 52.7% ± 0.12% of trials (in blue; upper panel). History-congruent perceptual choices were more frequent when perception was stimulus-incongruent (i.e., on *error* trials; lower panel), indicating that history effects impair performance in randomized psychophysical designs. B. Relative to randomly permuted data, we found highly significant autocorrelations of stimulus-congruence and history-congruence (dots indicate intercepts = 0 in trial-wise linear mixed effects modeling at p < 0.05). Across trials, the autocorrelation coefficients were best fit by an exponential function (adjusted *R*^2^ for stimulus-congruence: 0.53; history-congruence: 0.71) as compared to a linear function (adjusted *R*^2^ for stimulus-congruence: 0.52; history-congruence: 0.49). C. Here, we depict the number of consecutive trials at which autocorrelation coefficients exceeded the respective autocorrelation of randomly permuted data within individual partici­pants. For stimulus-congruence (upper panel), the lag of positive autocorrelation amounted to 3.24 ± 2.39 × 10^-3^ on average, showing a peak at trial t+1 after the index trial. For history-congruence (lower panel), the lag of positive autocorrelation amounted to 4.87 ± 3.36 × 10^-3^ on average, peaking at trial t+2 after the index trial. D. The smoothed probabilities of stimulus- and history-congruence (sliding windows of ±5 trials) fluctuated as *1/f noise*, i.e., at power densities that were inversely proportional to the frequency. E. The distribution of phase shift between fluctuations in stimulus- and history-congruence peaked at half a cycle (*π* denoted by dotted line). F. The average squared coherence between fluctuations in stimulus- and history-congruence (black dottet line) amounted to 6.49 ± 2.07 × 10^-3^% G. We observed faster response times (RTs) for both stimulus-congruence (as opposed to stimulus-incongruence, *β* = −0.14 ± 1.61 × 10^-3^, T(1.99 × 10^6^) = −85.91, p < 2.2 × 10^-308^) and history-congruence (*β* = −9.73 × 10^-3^ ± 1.38 × 10^-3^, T(1.99 × 10^6^) = −7.06, p = 1.66 × 10^-12^). H. The mode of perceptual processing (i.e., the difference between the smoothed probability of stimulus-vs. history-congruence) showed a quadratic relationship to RTs, with faster response times for stronger biases toward both external sensory information and internal predictions provided by perceptual history (*β*2 = −19.86 ± 0.52, T(1.98 × 10^6^) = −38.43, p = 5 × 10^-323^). The horizontal and vertical dotted lines indicate maximum RT and the associated mode, respectively. I. Confidence was enhanced for both stimulus-congruence (as opposed to stimulus-incongruence, *β* = 0.48 ± 1.38 × 10^-3^, T(2.06 × 10^6^) = 351.89, p < 2.2 × 10^-308^) and history-congruence (*β* = 0.04 ± 1.18 × 10^-3^, T(2.06 × 10^6^) = 36.86, p = 2.93 × 10^-297^). J. In analogy to RTs, we found a quadratic relationship between the mode of perceptual processing and confidence, which increased when both externally- and internally-biased modes grew stronger (*β*2 = 39.3 ± 0.94, T(2.06 × 10^6^) = 41.95, p < 2.2 × 10^-308^). The horizontal and vertical dotted lines indicate minimum confidence and the associated mode, respectively.

In addition, stimulus-congruence was significantly autocorrelated not only at the group-level, but also in individual participants, where the autocorrelation of stimulus-congruent perception exceeded the respective autocorrelation of randomly permuted data within an interval of 3.24 ± 2.39 × 10^-3^ trials (Figure 2C). In other words, if a participant’s experience was congruent (or incongruent) with the external stimulus information at a given trial, her perception was more likely to be stimulus-congruent (or incongruent) for approximately 3 trials into the future.

To further corroborate the autocorrelation of stimulus-congruence, we used logistic regression models that predicted the stimulus-congruence of perception at the index trial *t* = 0 from the stimulus-congruence at the preceding trials within a lag of 25 trials. We found that regression weights were significantly greater than zero for up to 16 trials (Supplemental Figure S3).

These results confirm that the ability to process sensory signals is not constant over time, but unfolds in multi-trial epochs of enhanced and reduced sensitivity to external information^8^. As a consequence of this autocorrelation, the dynamic probability of stimulus-congruent perception (i.e., computed in sliding windows of ± 5 trials; Figure 1C) fluctuated considerably within participants (average minimum: 35.46% ± 0.22%, maximum: 98.27% ± 0.07%). In line with previous findings^9^, such fluctuations in the sensitivity to external information had a power density that was inversely proportional to the frequency in the infra-slow spectrum^11^ (power ∼ 1/*f^β^*, *β* = −1.32 ± 3.14 × 10^-3^, T(1.84 × 10^5^) = −419.48, p < 2.2 × 10^-308^; Figure 2D). This feature, which is also known as *1/f noise* ^36, 37^, represents a characteristic of ongoing fluctuations in complex dynamic systems such as the brain^38^ and the cognitive processes it _entertains_^9,10, 13, 39, 40^.

### 5.2 Human perception fluctuates between external and internal modes of sensory processing

In a second step, we sought to explain why perception cycles through periods of enhanced and reduced sensitivity to external information^4,5^. We reasoned that observers may intermittently rely more strongly on internal information, i.e., on predictions about the environment that are constructed from previous experiences^19, 31^.

In perception, *serial dependencies* represent one of the most basic internal predictions that cause perceptual decisions to be systematically biased toward preceding choices^22–33^. Such effects of perceptual history mirror the continuity of the external world, in which the recent past often predicts the near future^30, 31, 34, 35, 41^. Therefore, as a metric for the perceptual impact of internal information, we computed whether the participant’s response at a given trial matched or differed from her response at the preceding trial (*history-congruent* and *history-incongruent perception*, respectively; Figure 1B and C).

First, we ensured that perceptual history played a significant role in perception despite the ongoing stream of external information. With a global average of 52.7% ± 0.12% history-congruent trials, we found a small but highly significant perceptual bias towards preceding experiences (*β* = 16.18 ± 1.07, T(1.09 × 10^3^) = 15.07, p = 10^-46^; Figure 2A) that was largely consistent across studies (Supplemental Figure 1B) and more pronounced in participants who were less sensitive to external sensory information (Supplemental Figure 1C). Logistic regression confirmed the internal information provided by perceptual history made a significant contribution to perception (*β* = 0.11 ± 5.79 × 10^-3^, z = 18.53, p = 1.1 × 10^-76^) over and above the ongoing stream of external sensory information (*β* = 2.2 ± 5.87 × 10^-3^, z = 375.11, p < 2.2 × 10^-308^) and general response biases toward one of the two potential outcomes (*β* = 15.19 ± 0.08, z = 184.98, p < 2.2 × 10^-308^; see Supplemental Figure S4A for model comparisons within individual participants).

In addition, we confirmed that history-congruence was not a corollary of the sequence of presented stimuli: History-congruent perceptual choices were more frequent at trials when perception was stimulus-incongruent (56.03% ± 0.2%) as opposed to stimulus-congruent (51.77% ± 0.11%, *β* = −4.26 ± 0.21, T(8.57 × 10^3^) = −20.36, p = 5.28 × 10^-90^; Figure 2A, lower panel). Despite being adaptive in auto-correlated real-world environments^19, 34, 35, 42^, perceptual history thus represented a source of error in the randomized experimental designs studied here^24, 28, 30, 31, 43^.

Second, we asked whether perception cycles through multi-trial epochs during which perception is characterized by stronger or weaker biases toward preceding experiences. Indeed, in close analogy to stimulus-congruence, history-congruence was significantly autocorrelated for up to 21 trials (Figure 2B). Following a peak at the first trial, the respective autocorrelation coefficients decreased exponentially over time (rate *γ* = −6.11 × 10^-3^ ± 5.69 × 10^-4^, T(6.75 × 10^4^) = −10.74, p = 7.18 × 10^-27^). History-congruence remained significantly autocorrelated when controlling for task difficulty (Supplemental Figure S2A) and the sequence of presented stimuli (Supplemental Figure S2B). In individual participants, the autocorrelation of history-congruence was elevated above randomly permuted data for a lag of 4.87 ± 3.36 × 10^-3^ trials (Figure 2C), confirming that the autocorrelation of history-congruence was not only a group-level phenomenon. The autocorrelation of history-congruence was confirmed by logistic regression models that successfully predicted the history-congruence of perception at an index trial *t* = 0 from the history-congruence at the preceding trials within a lag of 17 trials (Supplemental Figure S3).

Third, we asked whether the impact of internal information fluctuates as 1/f noise (i.e., a noise characteristic classically associated with fluctuations in the sensitivity to external information^9,10, 13, 39, 40^). The dynamic probability of history-congruent perception (i.e., computed in sliding windows of ± 5 trials; Figure 1C) varied considerably over time, ranging between a minimum of 12.77% ± 0.14% and a maximum 92.23% ± 0.14%. In analogy to stimulus-congruence, we found that history-congruence fluctuated as 1/f noise, with power densities that were inversely proportional to the frequency in the infra-slow spectrum^11^ (power ∼ 1/*f^β^*, *β* = −1.34 ± 3.16 × 10^-3^, T(1.84 × 10^5^) = −423.91, p < 2.2 × 10^-308^; Figure 2D). Finally, we ensured that fluctuations in stimulus- and history-congruence are linked to each other. When perceptual choices were less biased toward external information, participants relied more strongly on internal information acquired from perceptual history (and vice versa, *β* = −0.1 ± 8.59 × 10^-4^, T(2.1 × 10^6^) = −110.96, p < 2.2 × 10^-308^). Thus, while sharing the characteristic of 1/f noise, fluctuations in stimulus- and history-congruence were shifted against each other by approximately half a cycle and showed a squared coherence of 6.49 ± 2.07 × 10^-3^ % (Figure 2E and F; we report the average phase and coherence for frequencies below 0.1 1*/Ntrials*; see Methods for details).

In sum, our analyses indicate that perceptual decisions may result from a competition between external sensory signals with internal predictions provided by perceptual history. Crucially, we show that the impact of these external and internal sources of information is not stable over time, but fluctuates systematically, emitting overlapping autocorrelation curves and antiphase 1/f noise profiles.

These links between stimulus- and history-congruence suggest that the fluctuations in the impact of external and internal information may be generated by a unifying mechanism that causes perception to alternate between two opposing *modes* ^18^ (Figure 1D): During *external mode*, perception is more strongly driven by the available external stimulus information. Conversely, during *internal mode*, participants rely more heavily on internal predictions that are implicitly provided by preceding perceptual experiences. Fluctuations in mode (i.e., the degree of bias toward external versus internal information) may thus provide a novel explanation for ongoing fluctuations in the sensitivity to external information^4,5,18^.

### 5.3 Internal and external modes of processing facilitate response behavior and enhance confidence in human perceptual decision­making

Alternatively, however, fluctuating biases toward externally- and internally-oriented modes may not represent a perceptual phenomenon, but result from cognitive processes that are situated up- or downstream of perception. For instance, it may be argued that participants may be prone to stereotypically repeat the preceding choice when not attending to the experimental task. Thus, fluctuations in mode may arise due to systematic changes in the level of tonic arousal^44^ or on-task attention^45, 46^. Since arousal and attention typically link closely with response times^45, 47^ (RTs), this alternative explanation entails that RTs increase monotonically as one moves away from externally-biased and toward internally-biases modes of sensory processing.

As expected, stimulus-congruent (as opposed to stimulus-incongruent) choices were associated with faster responses (*β* = −0.14 ± 1.61 × 10^-3^, T(1.99 × 10^6^) = −85.91, p < 2.2 × 10^-308^; Figure 2G). Intriguingly, whilst controlling for the effect of stimulus-congruence, we found that history-congruent (as opposed to history-incongruent) choices were also characterized by shorter RTs (*β* = −9.73 × 10^-3^ ± 1.38 × 10^-3^, T(1.99 × 10^6^) = −7.06, p = 1.66 × 10^-12^; Figure 2G).

When analyzing the speed of response against the mode of sensory processing (Figure 2H), we found that RTs were shorter during externally-oriented perception (*β*1 = −11.07 ± 0.55, T(1.98 × 10^6^) = −20.14, p = 3.17 × 10^-90^). Crucially, as indicated by a quadratic relationship between the mode of sensory processing and RTs (*β*2 = −19.86 ± 0.52, T(1.98 × 10^6^) = −38.43, p = 5 × 10^-3^^23^), participants became faster at indicating their perceptual decision when biases toward both internal and external mode grew stronger. This argued against the view that the dynamics of pre-perceptual variables such as arousal or attention provide a plausible alternative explanation for the fluctuating perceptual impact of internal and external information.

Second, it may be assumed that participants tend to repeat preceding choices when they are not yet familiar with the experimental task, leading to history-congruent choices that are caused by insufficient training. In the Confidence database^20^, training effects were visible from RTs that were shortened by increasing exposure to the task (*β* = −7.53 × 10^-5^ ± 6.32 × 10^-7^, T(1.81 × 10^6^) = −119.15, p < 2.2 × 10^-308^). Intriguingly, however, history-congruent choices became more frequent with increased exposure to the task (*β* = 3.6 × 10^-5^ ± 2.54 × 10^-6^, z = 14.19, p = 10^-45^), speaking against the proposition that insufficient training induces seriality in response behavior.

As a third caveat, it could be argued that biases toward internal information reflect a post­perceptual strategy that repeats preceding choices when the subjective confidence in the perceptual decision is low. According to this view, subjective confidence should increase monotonically as biases toward external mode become stronger.

Stimulus-congruent (as opposed to stimulus-incongruent) choices were associated with enhanced confidence (*β* = 0.04 ± 1.18×10^-3^, T(2.06×10^6^) = 36.86, p = 2.93×10^-297^; Figure 2I). Yet whilst controlling for the effect of stimulus-congruence, we found that history-congruence also increased confidence (*β* = 0.48 ± 1.38 × 10^-3^, T(2.06 × 10^6^) = 351.89, p < 2.2 × 10^-308^; Figure 2I).

When depicted against the mode of sensory processing (Figure 2J), subjective confidence was indeed enhanced when perception was more externally-oriented (*β*_1_ = 92.63 ± 1, T(2.06 × 10^6^) = 92.89, p < 2.2 × 10^-308^). Importantly, however, participants were more confident in their perceptual decision for stronger biases toward both internal and external mode (*β*2 = 39.3 ± 0.94, T(2.06 × 10^6^) = 41.95, p < 2.2 × 10^-308^). In analogy to RTs, subjective confidence thus showed a quadratic relationship to the mode of sensory processing (Figure 2J), contradicting the notion that biases toward internal mode may reflect a post-perceptual strategy employed in situations of low subjective confidence.

The above results indicate that reporting behavior and metacognition do not map linearly onto the mode of sensory processing, suggesting that slow fluctuations in the respective impact of external and internal information are most likely to affect perception at an early level of sensory analysis^48,49^. Such low-level processing may integrate perceptual history with external inputs into a decision variable^50^ that influences not only perceptual choices, but also downstream functions such as speed of response and subjective confidence. Consequently, our findings predict that human participants lack full metacognitive insight into how strongly external signals and internal predictions contribute to perceptual decision-making. Stronger biases toward perceptual history thus lead to two seemingly contradictory effects: more frequent errors (Supplemental Figure 1C) and increasing subjective confidence (Figure 2I-J).

This observation generates an intriguing prediction regarding the association of between­mode fluctuations and perceptual metacognition: Metacognitive efficiency should be lower in individuals who spend more time in internal mode, since their confidence reports are less predictive of whether the corresponding perceptual decision is correct. We computed each participant’s M-ratio^51^ (meta-d’/d’ = 0.85 ± 0.02) to probe this hypothesis independently of inter-individual differences in perceptual performance. Indeed, we found that biases toward internal information (i.e., as defined by the average probability of history-congruence) were stronger in participants with lower metacognitive efficiency (*β* = −2.98 × 10^-3^ ± 9.82 × 10^-4^, T(4.14 × 10^3^) = −3.03, p = 2.43 × 10^-3^).

### 5.4 Fluctuations between internal and external mode modulate perceptual performance beyond the effect of general response biases

The above sections provide correlative evidence that recurring intervals of stronger perceptual history temporally reduce the participants’ sensitivity to external information. Importantly, the history-dependent biases that characterize internal mode processing must be differentiated from general response biases. In binary perceptual decision-making, general response biases are defined by a propensity to choose one of the two outcomes more often than the alternative. Indeed, in the experiments considered here, participants selected the more frequent of the two possible outcomes in 58.71% ± 0.22% of trials.

Two caveats have to be considered to make sure that the effect of history-congruence is distinct from the effect of general response biases. First, history-congruent states become more likely for larger response biases that cause a increasing imbalance in the likelihood of the two outcomes (*β* = 0.24 ± 6.93 × 10^-4^, T(2.09 × 10^6^) = 342.43, p < 2.2 × 10^-308^). One may thus ask whether the autocorrelation of history-congruence could be entirely driven by general response biases. Yet the above analyses account for general response biases by computing group-level autocorrelations (see Figure 2C) relative to randomly permuted data (i.e., by subtracting the autocorrelation of randomly permuted data from the raw autocorrelation curve). This precludes that general response biases contribute to the observed autocorrelation of history-congruence (see Supplemental Figure S5 for a visualization of the correction procedure for simulated data with general response biases ranging from 60 to 90%).

Second, it may be argued that fluctuations in perceptual performance may be solely driven by ongoing changes in the strength of general response biases. To assess the links between dynamic fluctuations in stimulus-congruence on the one hand and history-congruence as well as general response bias on the other hand, we computed all variables as dynamic probabilities in sliding windows of ± 5 trials (see Figure 1C). Linear mixed effects modeling indicated that fluctuations in history-congruent biases were larger in amplitude than the corresponding fluctuations in general response biases (*β*_0_ = 0.03 ± 7.34 × 10^-3^, T(64.94) = 4.46, p = 3.28 × 10^-5^). Crucially, ongoing fluctuations in history-congruence had a significant effect on stimulus-congruence (*β*1 = −0.05 ± 5.63 × 10^-4^, T(2.1 × 10^6^) = −84.21, p < 2.2 × 10^-308^) beyond the effect of ongoing changes in general response biases (*β*2 = −0.06 ± 5.82 × 10^-4^, T(2.1 × 10^6^) = −103.51, p < 2.2 × 10^-308^). In sum, the above control analyses confirm that the observed influence of preceding choices on perceptual decision-making cannot not be reduced to general response biases.

### 5.5 Internal mode is characterized by lower thresholds as well as by history-dependent changes in biases and lapses

In a final control analysis, we asked whether history-independent changes in biases and lapses may provide an alternative explanation of internal mode processing. To this end, we estimated full and history-conditioned psychometric curves to investigate how internal and external mode relate to biases (i.e., the horizontal position of the psychometric curve), lapses (i.e, the asymptotes of the psychometric curve) and thresholds (i.e., 1/sensitivity, estimated from the slope of the psychometric curve). We used a maximum likelihood procedure to predict trial-wise choices *y* (*y* = 0 and *y* = 1 for outcomes A and B respectively) from the choice probabilities *yp*. *yp* was computed from difficulty-weighted inputs *sw* via a parametric error function defined by the parameters *γ* (lower lapse), *δ* (upper lapse), *µ* (bias) and *t* (threshold; see Methods for details):

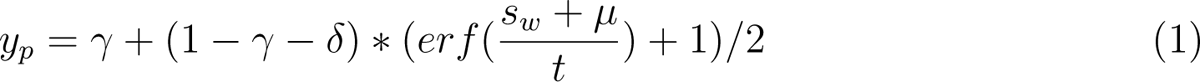

Across the full dataset (i.e., irrespective of the preceding perceptual choice *yt*-1), biases *µ* were distributed around zero (−0.05 ± 0.03; *β*_0_ = 7.37 × 10^-3^ ± 0.09, T(36.8) = 0.08, p = 0.94; see Figure 3A and B, upper panel). When conditioned on perceptual history, biases *µ* varied according to the preceding perceptual choice, with negative biases for *Yt*-1 = 0 (−0.22 ± 0.04; *β*_0_ = 0.56 ± 0.12, T(43.39) = 4.6, p = 3.64 × 10^-5^) and positive biases for *Yt*-1 = 1 (0.29 ± 0.03; *β*_0_ = 0.56 ± 0.12, T(43.39) = 4.6, p = 3.64 × 10^-5^). Absolute biases |*µ*| were larger in internal mode (1.84 ± 0.03) as compared to external mode (0.86 ± 0.02; *β*_0_ = −0.62 ± 0.07, T(45.62) = −8.38, p = 8.59 × 10^-11^; controlling for differences in lapses and thresholds). Lower and upper lapses amounted to *γ* = 0.13 ± 2.83 × 10^-3^ and *δ* = 0.1 ± 2.45 × 10^-3^ (see Figure 3A, C and D). Lapses were larger in internal mode (*γ* = 0.17 ± 3.52 × 10^-3^, *δ* = 0.14 ± 3.18 × 10^-3^) as compared to external mode (*γ* = 0.1 ± 2.2 × 10^-3^, *δ* = 0.08 ± 2 × 10^-3^; *β*_0_ = −0.05 ± 5.73 × 10^-3^, T(47.03) = −9.11, p = 5.94 × 10^-12^; controlling for differences in biases and thresholds).

**Figure 3.**
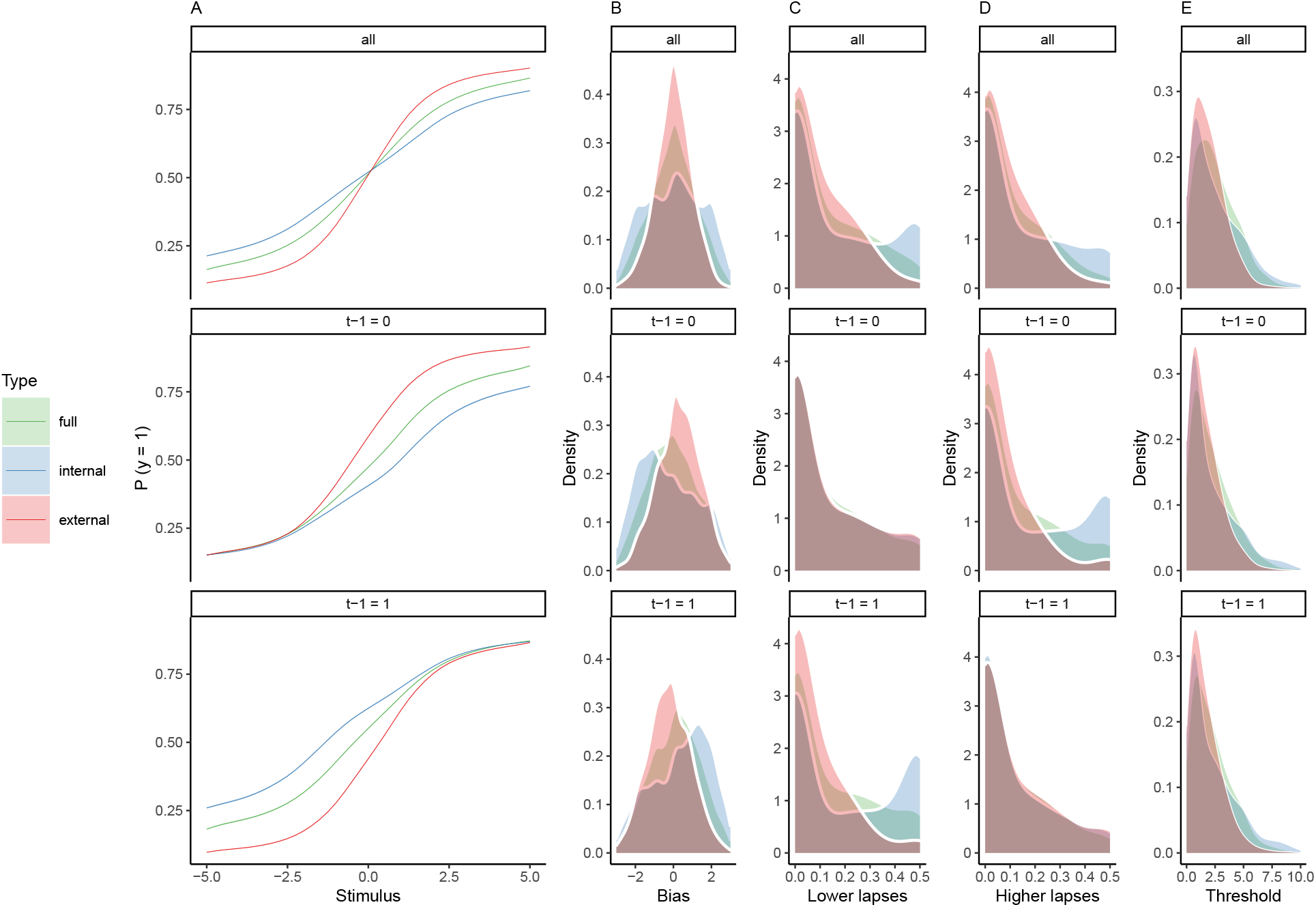
Full and history-conditioned psychometric functions across modes in humans. A. Here, we show average psychometric functions for the full dataset (upper panel) and conditioned on perceptual history (*yt*-1 = 1 and *yt*-1 = 0; middle and lower panel) across modes (green line) and for internal mode (blue line) and external mode (red line) separately. B. Across the full dataset, biases *µ* were distributed around zero (*β*_0_ = 7.37 × 10^-3^ ± 0.09, T(36.8) = 0.08, p = 0.94; upper panel), with larger absolute biases |*µ*| for internal as compared to external mode (*β*_0_ = −0.62 ± 0.07, T(45.62) = −8.38, p = 8.59 × 10^-11^; controlling for differences in lapses and thresholds). When conditioned on perceptual history, we observed negative biases for *yt*-1 = 0 (*β*_0_ = 0.56 ± 0.12, T(43.39) = 4.6, p = 3.64 × 10^-5^; middle panel) and positive biases for *yt*-1 = 1 (*β*_0_ = 0.56 ± 0.12, T(43.39) = 4.6, p = 3.64 × 10^-5^; lower panel). C. Lapse rates were higher in internal mode as compared to external mode (*β*_0_ = −0.05 ± 5.73 × 10^-3^, T(47.03) = −9.11, p = 5.94 × 10^-12^; controlling for differences in biases and thresholds; see upper panel and subplot D). Importantly, the between-mode difference in lapses depended on perceptual history: We found no significant difference in lower lapses *γ* for *yt*-1 = 0 (*β*_0_ = 0.01 ± 7.77 × 10^-3^, T(33.1) = 1.61, p = 0.12; middle panel), but a significant difference for *yt*-1 = 1 (*β*_0_ = −0.11 ± 0.01, T(40.11) = −9.59, p = 6.14 × 10^-12^; lower panel). D. Conversely, higher lapses *δ* were significantly increased for *yt*-1 = 0 (*β*_0_ = −0.1 ± 9.58 × 10^-3^, T(36.87) = −10.16, p = 3.06 × 10^-12^; middle panel), but not for *yt*-1 = 1 (*β*_0_ = 0.01 ± 7.74 × 10^-3^, T(33.66) = 1.58, p = 0.12; lower panel). E. The thresholds *t* were larger in internal as compared to external mode (*β*_0_ = −1.77 ± 0.25, T(50.45) = −7.14, p = 3.48 × 10^-9^; controlling for differences in biases and lapses) and were not modulated by perceptual history (*β*_0_ = 0.04 ± 0.06, T(2.97 × 10^3^) = 0.73, p = 0.47).

Conditioning on the previous perceptual choice revealed that the between-mode difference in lapse was not general, but depended on perceptual history: For *yt*-1 = 0, only higher lapses *δ* differed between internal and external mode (*β*_0_ = −0.1 ± 9.58 × 10^-3^, T(36.87) = −10.16, p = 3.06 × 10^-12^), whereas lower lapses *γ* did not (*β*_0_ = 0.01 ± 7.77 × 10^-3^, T(33.1) = 1.61, p = 0.12). Vice versa, for *yt*-1 = 1, lower lapses *γ* differed between internal and external mode (*β*_0_ = −0.11 ± 0.01, T(40.11) = −9.59, p = 6.14 × 10^-12^), whereas higher lapses *δ* did not (*β*_0_ = 0.01 ± 7.74 × 10^-3^, T(33.66) = 1.58, p = 0.12).

Thresholds *t* were estimated at 3 ± 0.06 (see Figure 3A and E). Thresholds *t* were larger in internal mode (3.66 ± 0.09) as compared to external mode (2.02 ± 0.03; *β*_0_ = −1.77 ± 0.25, T(50.45) = −7.14, p = 3.48 × 10^-9^; controlling for differences in biases and lapses). In contrast to the bias *µ* and the lapse rates *γ* and *δ*, thresholds *t* were not modulated by perceptual history (*β*_0_ = 0.04 ± 0.06, T(2.97 × 10^3^) = 0.73, p = 0.47).

In sum, the above analyses showed that internal and external mode differ with respect to biases, lapses and thresholds. Internally-biased processing was characterized by higher thresholds, indicating a reduced sensitivity to sensory information, as well as by larger biases and lapses. Importantly, between-mode differences in biases and lapses strongly depended on perceptual history. This confirmed that internal mode processing cannot be explained solely on the ground of a general (i.e., history-independent) increase in lapses or bias.

### 5.6 Mice waver between external and internal modes of perceptual decision-making

In a prominent functional explanation for serial dependencies^22–28, 32, 33, 48^, perceptual history is cast as an internal prediction that leverages the temporal autocorrelation of natural environments for efficient decision-making^30, 31, 34, 35, 41^. We reasoned that, since this autocorrelation is one of the most basic features of our sensory world, fluctuating biases toward preceding perceptual choices should not be a uniquely human phenomenon.

To test whether externally and internally oriented modes of processing exist beyond the human mind, we analyzed data on perceptual decision-making in mice that were extracted from the International Brain Laboratory (IBL) dataset^21^. Here, we restricted our analyses to the *basic* task^21^, in which mice responded to gratings of varying contrast that appeared either in the left or right hemifield of with equal probability. We excluded sessions in which mice did not respond correctly to stimuli presented at a contrast above 50% in more than 80% of trials (see Methods), which yielded a final sample of N = 165 adequately trained mice that went through 1.46 million trials.

In line with humans, mice were biased toward perceptual history in 54.03% ± 0.17% of trials (T(164) = 23.65, p = 9.98 × 10^-55^; Figure 4A and Supplemental Figure S1D). Perceptual history effects remained significant (*β* = 0.51 ± 4.49 × 10^-3^, z = 112.84, p = 0) when controlling for external sensory information (*β* = 2.96 ± 4.58 × 10^-3^, z = 646.1, p = 0) and general response biases toward one of the two potential outcomes (*β* = −1.78 ± 0.02, z = −80.64, p < 2.2 × 10^-308^; see Supplemental Figure S4C-D for model comparisons and *β* values computed within individual mice).

**Figure 4.**
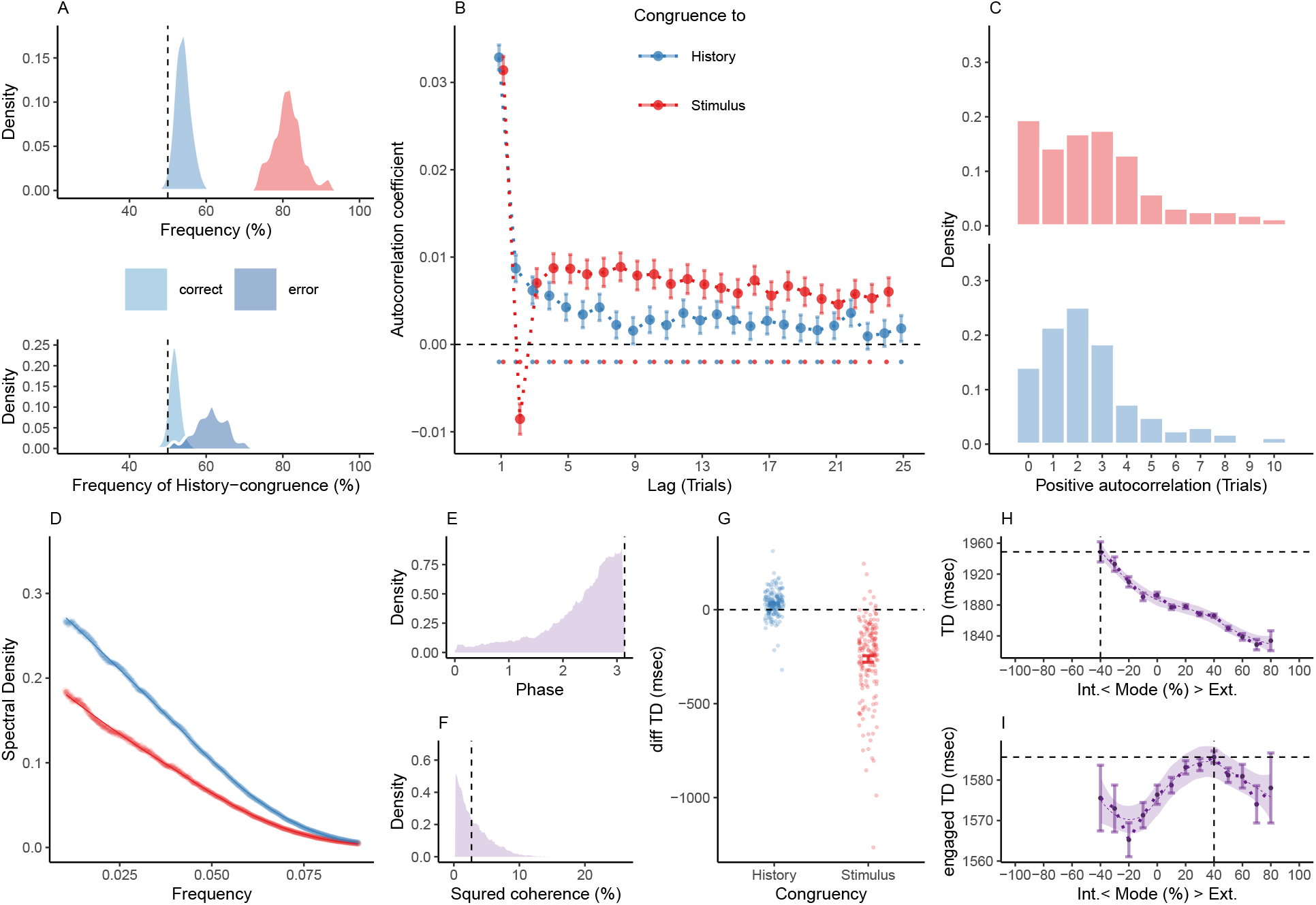
Internal and external modes in murine perceptual decision-making. A. In mice, 81.37% ± 0.3% of trials were stimulus-congruent (in red) and 54.03% ± 0.17% of trials were history-congruent (in blue; upper panel). History-congruent perceptual choices were not a consequence of the experimental design, but a source of error, as they were more frequent on stimulus-incongruent trials (lower panel). B. Relative to randomly permuted data, we found highly significant autocorrelations of stimulus-congruence and history-congruence (dots indicate intercepts = 0 in trial-wise linear mixed effects modeling at p < 0.05). Please note that the negative autocorrelation of stimulus-congruence at trial 2 was a consequence of the experimental design (see Supplemental Figure 2D-F). As in humans, autocorrelation coefficients were best fit by an exponential function (adjusted *R*^2^ for stimulus-congruence: 0.44; history-congruence: 0.52) as compared to a linear function (adjusted *R*^2^ for stimulus-congruence: 3.16 × 10^-3^; history-congruence: 0.26). C. For stimulus-congruence (upper panel), the lag of positive autocorrelation was longer in comparison to humans (4.59 ± 0.06 on average). For history-congruence (lower panel), the lag of positive autocorrelation was slightly shorter relative to humans (2.58 ± 0.01 on average, peaking at trial t+2 after the index trial). D. In mice, the dynamic probabilities of stimulus- and history-congruence (sliding windows of ±5 trials) fluctuated as *1/f noise*. E. The distribution of phase shift between fluctuations in stimulus- and history-congruence peaked at half a cycle (*π* denoted by dotted line). F. The average squared coherence between fluctuations in stimulus- and history-congruence (black dottet line) amounted to 3.45 ± 0.01% G. We observed shorter trial durations (TDs) for stimulus-congruence (as opposed to stimulus-incongruence, *β* = −1.12 ± 8.53 × 10^-3^, T(1.34 × 10^6^) = −131.78, p < 2.2 × 10^-308^), but longer TDs for history-congruence (*β* = 0.06 ± 6.76 × 10^-3^, T(1.34 × 10^6^) = 8.52, p = 1.58 × 10^-17^). H. TDs decreased monotonically for stronger biases toward external mode (*β*_1_ = −4.16 × 10^4^ ± 1.29 × 10^3^, T(1.35 × 10^6^) = −32.31, p = 6.03 × 10^-229^). The horizontal and vertical dotted lines indicate maximum TD and the associated mode, respectively. I. For TDs that differed from the median TD by no more than 1.5 x MAD (median absolute distance^52^), mice exhibited a quadratic component in the relationship between the mode of sensory processing and TDs (*β*2 = −1.97 × 10^3^ ± 843.74, T(1.19 × 10^6^) = −2.34, p = 0.02, Figure 4I). This explorative post-hoc analysis focuses on trials at which mice engage more swiftly with the experimental task. The horizontal and vertical dotted lines indicate maximum TD and the associated mode, respectively.

In the *basic* task of the IBL dataset^21^, stimuli were presented at random in either the left or right hemifield. Stronger biases toward perceptual history should therefore decrease perceptual performance. Indeed, history-congruent choices were more frequent when perception was stimulus-incongruent (61.59% ± 0.07%) as opposed to stimulus-congruent (51.81% ± 0.02%, T(164) = 31.37, p = 3.36 × 10^-71^; T(164) = 31.37, p = 3.36 × 10^-71^; Figure 4A, lower panel), confirming that perceptual history was a source of error^24,28, 30, 31, 43^ as opposed to a feature of the experimental paradigm. Overall, perception was stimulus-congruent in 81.37% ± 0.3% of trials (Figure 4A).

At the group level, we found significant autocorrelations in both stimulus-congruence (86 consecutive trials) and history-congruence (8 consecutive trials), which remained significant when taking into account the respective autocorrelation of task difficulty and external stimulation (Supplemental Figure 2C-D). In contrast to humans, mice showed a negative autocorrelation coefficient of stimulus-congruence at trial 2. This was due to a feature of the experimental design: Errors at a contrast above 50% were followed by a high-contrast stimulus at the same location. Thus, stimulus-incongruent choices on easy trials were more likely to be followed by stimulus-congruent perceptual choices that were facilitated by high-contrast visual stimuli^21^.

The autocorrelation of history-congruence closely overlapped with the human data and decayed exponentially after a peak at the first trial (rate *γ* = −6.7 × 10^-3^ ± 5.94 × 10^-4^, T(3.69 × 10^4^) = −11.27, p = 2.07 × 10^-29^; Figure 4B). On the level of individual mice, autocorrelation coefficients were elevated above randomly permuted data within a lag of 4.59 ± 0.06 trials for stimulus-congruence and 2.58 ± 0.01 trials for history-congruence (Figure 4C).

To further corroborate a significant autocorrelation of stimulus- and history-congruence in mice, we used logistic regression models that predicted the stimulus-/history-congruence of perception at the index trial *t* = 0 from the stimulus/history-congruence at the preceding trials within a lag of 25 trials. We found that regression weights were significantly greater than zero for more than 25 trials for stimulus-congruence. For history-congruence, regression weights significantly greater than zero for 8 trials prior to the index trial (Supplemental Figure S3). In analogy to humans, mice showed anti-phase 1/f fluctuations in the sensitivity to internal and external information (Figure 4D-F).

Next, we asked how external and internal modes relate to the trial duration (TD, a coarse measure of RT in mice that spans the interval from stimulus onset to feedback^21^). Stimulus-congruent (as opposed to stimulus-incongruent) choices were associated with shorter TDs (*δ* = −262.48 ± 17.1, T(164) = −15.35, p = 1.55 × 10^-33^), while history-congruent choices were characterized by longer TDs (*δ* = 30.47 ± 5.57, T(164) = 5.47, p = 1.66 × 10^-7^; Figure 4G). Across the full spectrum of the available data, TDs showed a linear relationship with the mode of sensory processing, with shorter TDs during external mode (*β*_1_ = −4.16 × 10^4^ ± 1.29 × 10^3^, T(1.35 × 10^6^) = −32.31, p = 6.03 × 10^-229^, Figure 4H). However, an explorative post-hoc analysis limited to TDs that differed from the median TD by no more than 1.5 x MAD (median absolute distance^52^) indicated that, when mice engaged with the task more swiftly, TDs did indeed show a quadratic relationship with the mode of sensory processing (*β*2 = −1.97 × 10^3^ ± 843.74, T(1.19 × 10^6^) = −2.34, p = 0.02, Figure 4I).

As in humans, it is an important caveat to consider whether the observed serial dependencies in murine perception reflect a phenomenon of perceptual inference, or, alternatively, an unspecific strategy that occurs at the level of reporting behavior. We reasoned that, if mice indeed tended to repeat previous choices as a general response pattern, history effects should decrease during training of the perceptual task. We therefore analyzed how stimulus- and history-congruent perceptual choices evolved across sessions in mice that, by the end of training, achieved proficiency (i.e., stimulus-congruence ≥ 80%) in the *basic* task of the IBL dataset^21^.

As expected, we found that stimulus-congruent perceptual choices became more frequent (*β* = 0.34 ± 7.13 × 10^-3^, T(8.51 × 10^3^) = 47.66, p < 2.2 × 10^-308^; Supplemental Figure S6) and TDs were progressively shortened (*β* = −22.14 ± 17.06, T(1.14 × 10^3^) = −1.3, p < 2.2 × 10^-308^) across sessions. Crucially, the frequency of history-congruent perceptual choices also increased during training (*β* = 0.13 ± 4.67 × 10^-3^, T(8.4 × 10^3^) = 27.04, p = 1.96 × 10^-154^; Supplemental Figure S6).

As in humans, longer within-session task exposure was associated with an increase in history-congruence (*β* = 3.6 × 10^-5^ ± 2.54 × 10^-6^, z = 14.19, p = 10^-45^) and a decrease in TDs (*β* = −0.1 ± 3.96 × 10^-3^, T(1.34 × 10^6^) = −24.99, p = 9.45 × 10^-138^). In sum, these findings strongly argue against the proposition that mice show biases toward perceptual history due to an unspecific response strategy.

As in humans, fluctuations in the strength of history-congruent biases were, (i), larger in amplitude than the corresponding fluctuations in general response biases (*β*_0_ = −5.26 × 10^-3^ ± 4.67 × 10^-4^, T(2.12 × 10^3^) = −11.28, p = 1.02 × 10^-28^) and, (ii), had a significant effect on stimulus-congruence (*β*1 = −0.12 ± 7.17 × 10^-4^, T(1.34 × 10^6^) = −168.39, p < 2.2 × 10^-308^) beyond the effect of ongoing changes in general response biases (*β*2 = −0.03 ± 6.94 × 10^-4^, T(1.34 × 10^6^) = −48.14, p < 2.2 × 10^-308^). This confirmed that, in both humans and mice, perceptual performance is modulated by systematic fluctuations between externally- and internally-oriented modes of sensory processing.

Finally, we fitted full and history-conditioned psychometric curves to the data from the IBL database. When estimated based on the full dataset (i.e., irrespective of the preceding perceptual choice *yt*-1), biases *µ* were distributed around zero (3.87 × 10^-3^ ± 9.81 × 10^-3^; T(164) = 0.39, p = 0.69; Figure 5A and B, upper panel). When conditioned on the preceding perceptual choice, biases were negative for *yt*-1 = 0 (−0.02 ± 8.7 × 10^-3^; T(164) = −1.99, p = 0.05; Figure 5A and B, middle panel) and positive for *yt*-1 = 1 (0.02 ± 9.63 × 10^-3^; T(164) = 1.91, p = 0.06; Figure 5A and B, lower panel). As in humans, mice showed larger biases during internal mode (0.14 ± 7.96 × 10^-3^) as compared to external mode (0.07 ± 8.7 × 10^-3^; *β*_0_ = −0.18 ± 0.03, T = −6.38, p = 1.77 × 10^-9^; controlling for differences in lapses and thresholds).

**Figure 5.**
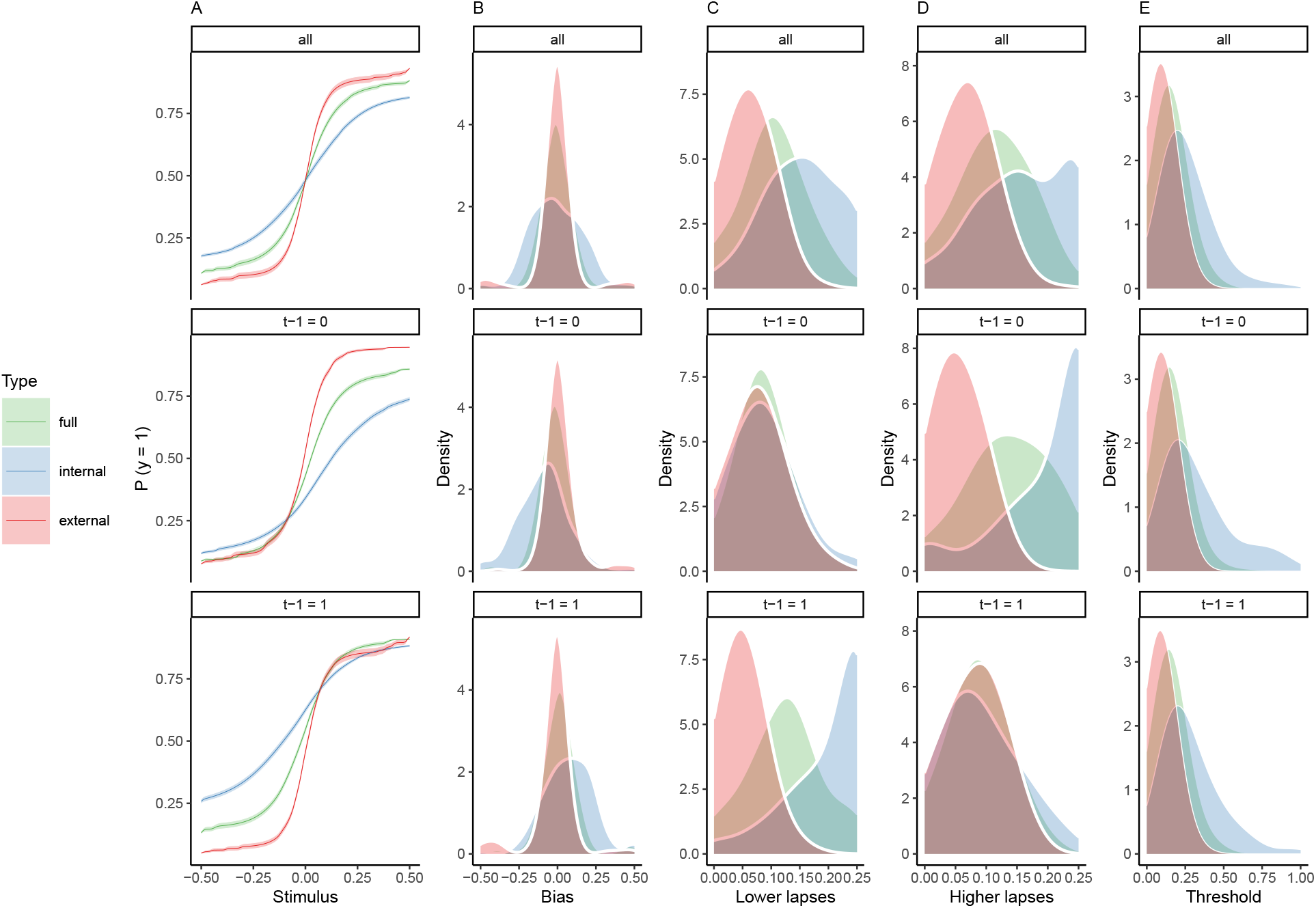
Full and history-conditioned psychometric functions across modes in mic. A. Here, we show average psychometric functions for the full IBL dataset (upper panel) and conditioned on perceptual history (*yt*-1 = 1 and *yt*-1 = 0; middle and lower panel) across modes (green line) and for internal mode (blue line) and external mode (red line) separately. B. Across the full dataset, biases *µ* were distributed around zero (T(164) = 0.39, p = 0.69; upper panel), with larger absolute biases |*µ*| for internal as compared to external mode (*β*_0_ = −0.18 ± 0.03, T = −6.38, p = 1.77 × 10^-9^; controlling for differences in lapses and thresholds). When conditioned on perceptual history, we observed negative biases for *yt*-1 = 0 (T(164) = −1.99, p = 0.05; middle panel) and positive biases for *yt*-1 = 1 (T(164) = 1.91, p = 0.06; lower panel). C. Lapse rates were higher in internal as compared to external mode (*β*_0_ = −0.11 ± 4.39 × 10^-3^, T = −24.8, p = 4.91 × 10^-57^; controlling for differences in biases and thresholds; upper panel, see also subplot D). For *yt*-1 = 1, the difference between internal and external mode was more pronounced for lower lapses *γ* (T(164) = −18.24, p = 2.68 × 10^-41^) as compared to higher lapses *δ* (see subplot D). In mice, lower lapses *γ* were significantly elevated during internal mode irrespective of the preceding perceptual choice (middle panel: lower lapses *γ* for *yt*-1 = 0; T(164) = −2.5, p = 0.01, lower panel: lower lapses *γ* for *yt*-1 = 1; T(164) = −32.44, p = 2.92 × 10^-73^). D. For *yt*-1 = 0, the difference between internal and external mode was more pronounced for higher lapses *δ* (T(164) = 21.44, p = 1.93 × 10^-49^, see subplot C). Higher lapses were significantly elevated during internal mode irrespective of the preceding perceptual choice (middle panel: higher lapses *δ* for *yt*-1 = 0; T(164) = −28.29, p = 5.62 × 10^-65^ lower panel: higher lapses *δ* for *yt*-1 = 1; T(164) = −2.65, p = 8.91 × 10^-3^;). E. Thresholds *t* were higher in internal as compared to external mode (*β*_0_ = −0.28 ± 0.04, T = −7.26, p = 1.53 × 10^-11^; controlling for differences in biases and lapses) and were not modulated by perceptual history (T(164) = 0.94, p = 0.35).

Lower and upper lapses amounted to *γ* = 0.1 ± 4.35 × 10^-3^ and *δ* = 0.11 ± 4.65 × 10^-3^ (see Figure 5A, C and D). Lapse rates were higher in internal mode (*γ* = 0.15 ± 5.14 × 10^-3^, *δ* = 0.16 ± 5.79 × 10^-3^) as compared to external mode (*γ* = 0.06 ± 3.11 × 10^-3^, *δ* = 0.07 ± 3.34 × 10^-3^; *β*_0_ = −0.11 ± 4.39 × 10^-3^, T = −24.8, p = 4.91 × 10^-57^; controlling for differences in biases and thresholds).

For *yt*-1 = 0, the difference between internal and external mode was more pronounced for higher lapses *δ* (T(164) = 21.44, p = 1.93 × 10^-49^). Conversely, for *yt*-1 = 1, the difference between internal and external mode was more pronounced for lower lapses *γ* (T(164) = −18.24, p = 2.68 × 10^-41^). In contrast to the human data, higher lapses *δ* and lower lapses *γ* were significantly elevated during internal mode irrespective of the preceding perceptual choice (higher lapses *δ* for *yt*-1 = 1: T(164) = −2.65, p = 8.91 × 10^-3^; higher lapses *δ* for *yt*-1 = 0: T(164) = −28.29, p = 5.62 × 10^-65^; lower lapses *γ* for *yt*-1 =1: T(164) = −32.44, p = 2.92 × 10^-73^; lower lapses *γ* for *yt*-1 = 0: T(164) = −2.5, p = 0.01).

In mice, thresholds *t* amounted to 0.15 ± 6.52 × 10^-3^ (see Figure 5A and E) and were higher in internal mode (0.27 ± 0.01) as compared to external mode (0.09 ± 4.44 × 10^-3^; *β*_0_ = −0.28 ± 0.04, T = −7.26, p = 1.53 × 10^-11^; controlling for differences in biases and lapses). Thresholds *t* were not modulated by perceptual history (T(164) = 0.94, p = 0.35).

In sum, the above analyses of the psychometric function in mice corroborated our findings in humans. Higher thresholds indicated a reduced sensitivity to external information during internal mode. Additionally, internally-biased processing was characterized history-dependent modulation of biases and lapses.

### 5.7 Fluctuations in mode result from coordinated changes in the impact of external and internal information on perception

The empirical data presented above indicate that, for both humans and mice, perception fluctuates between internal an external modes, i.e., multi-trial epochs that are character­ized by enhanced sensitivity toward either internal or external information. Since natural environments typically show high temporal redundancy^34^, previous experiences are often good predictors of new stimuli^30, 31, 35, 41^. Serial dependencies may therefore induce autocorre­lations in perception by serving as an internal prediction (or *memory* processes^9,13^) about the environment that actively integrates noisy sensory information over time^53^.

Previous work has shown that such internal predictions are built by dynamically updating the estimated probability of being in a particular perceptual state from the sequence of preceding experiences^35, 48, 54^. The integration of sequential inputs may lead to accumulating effects of perceptual history that progressively override incoming sensory information, enabling internal mode processing^19^. However, since such a process would lead to internal biases that may eventually become impossible to overcome^55^, we assumed that changes in mode may additionally be driven by ongoing wave-like fluctuations^9,13^ in the perceptual impact of external and internal information that occur *irrespective* of the sequence of previous experiences and temporarily de-couple the decision variable from implicit internal representations of the environment^19^.

Following Bayes’ theorem, we reasoned that binary perceptual decisions depend on the posterior log ratio *L* of the two alternative states of the environment that participants learn about via noisy sensory information^54^. We computed the posterior by combining the sensory evidence available at time-point *t* (i.e., the log likelihood ratio *LLR*) with the prior probability *ψ*:

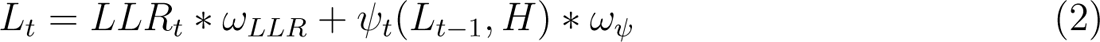

We derived the prior probability *ψ* at timepoint *t* from the posterior probability of perceptual outcomes at timepoint *Lt*-1. Since a switch between the two states can occur at any time, the effect of perceptual history varies according to both the sequence of preceding experiences and the estimated stability of the external environment (i.e., the *hazard rate H*^54^):

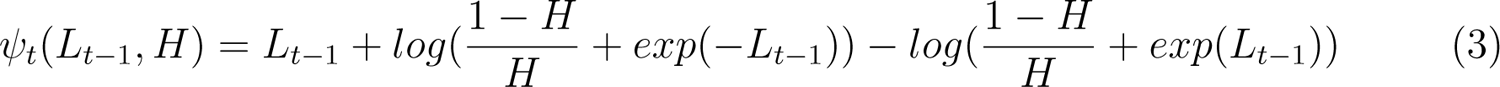

The *LLR* was computed from inputs *st* by applying a sigmoid function defined by parameter *α* that controls the sensitivity of perception to the available sensory information (see Methods for detailed equations on humans and mice):

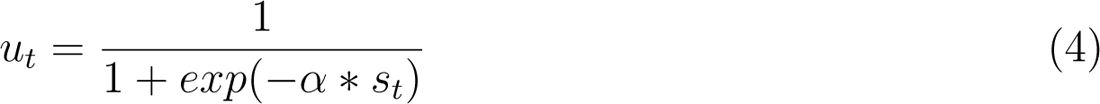

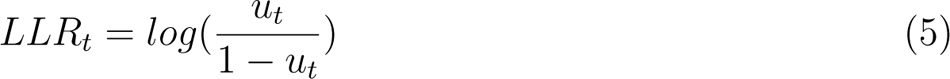

To allow for *bimodal inference*, i..e, alternating periods of internally- and externally-biased modes of perceptual processing that occur irrespective of the sequence of preceding experiences, we assumed that the relative influences of likelihood and prior show coherent anti-phase fluctuations governed by *ωLLR* and *ωψ* that are determined by amplitude *a*, frequency *f* and phase *p*:

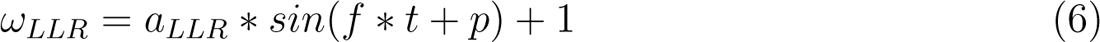

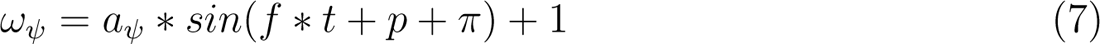

Finally, a sigmoid transform of the posterior *L_t_* yields the probability of observing the perceptual decision *yt* at a temperature determined by *ζ* ^-1^:

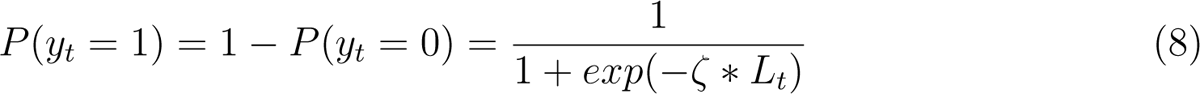

Fitting the bimodal inference model outlined above to behavioral data (see Methods for details) characterizes each subject by a sensitivity parameter *α* that captures how strongly perception is driven by the available sensory information, and a hazard rate parameter *H* that controls how heavily perception is biased by perceptual history. As a sanity check for model fit, we tested whether the frequency of stimulus- and history-congruent trials in the Confidence database^20^ and IBL database^21^ correlate with the estimated parameters *α* and *H*, respectively. As expected, the estimated sensitivity toward stimulus information *α* was positively correlated with the frequency of stimulus-congruent perceptual choices (humans: *β* = 8.4 ± 0.26, T(4.31 × 10^3^) = 32.87, p = 1.3 × 10^-2^^11^; mice: *β* = 1.93 ± 0.12, T(2.07 × 10^3^) = 16.21, p = 9.37 × 10^-56^). Likewise, *H* was negatively correlated with the frequency of history-congruent perceptual choices (humans: *β* = −11.84 ± 0.5, T(4.29 × 10^3^) = −23.5, p = 5.16 × 10^-1^^15^; mice: *β* = −6.18 ± 0.66, T(2.08 × 10^3^) = −9.37, p = 1.85 × 10^-20^).

Our behavioral analyses have shown that humans and mice showed significant effects of percep­tual history that impaired performance in randomized psychophysical experiments^24, 28, 30, 31, 43^ (Figure 2A and 3A). We therefore expected that humans and mice underestimated the true hazard rate *H* of the experimental environments (Confidence database^20^: *H_Humans_ =* 0.5 ± 1.58 × 10^-5^); IBL database^21^: *H_Mice_* = 0.49 ± 6.47 × 10^-5^). Indeed, when fitting the bimodal inference model outlined above to the trial-wise perceptual choices (see Methods), we found that the estimated (i.e., subjective) hazard rate *H* was lower than *H* for both humans (H = 0.45 ± 4.8× 10^-5^, *β* = −6.87 ± 0.94, T(61.87) = −7.33, p = 5.76× 10^-10^) and mice (H = 0.46 ± 2.97× 10^-4^, *β* = −2.91 ± 0.34, T(112.57) = −8.51, p = 8.65× 10^-14^).

Simulations from the bimodal inference model (based on the posterior model parameters obtained in humans; see Methods for details) closely matched the empirical results outlined above: Simulated perceptual decisions resulted from a competition of perceptual history with incoming sensory signals (Figure 6A). Stimulus- and history-congruence were significantly auto-correlated (Figure 6B-C), fluctuating in anti-phase as 1/f noise (Figure 6D-F). Simulated posterior certainty^28, 30, 50^ (i.e., the absolute of the posterior log ratio |*L_t_* |) showed a quadratic relationship to the mode of sensory processing (Figure 6H), mirroring the relation of RTs and confidence reports to external and internal biases in perception (Figure 2G-H and Figure 4G-H). Crucially, the overlap between empirical and simulated data broke down when we removed the anti-phase oscillations (*ωLLR* and/or *ωψ*) or the accumulation of evidence over time (i.e., setting *H* to 0.5) from the bimodal inference model (see Supplemental Figure S7-10).

**Figure 6.**
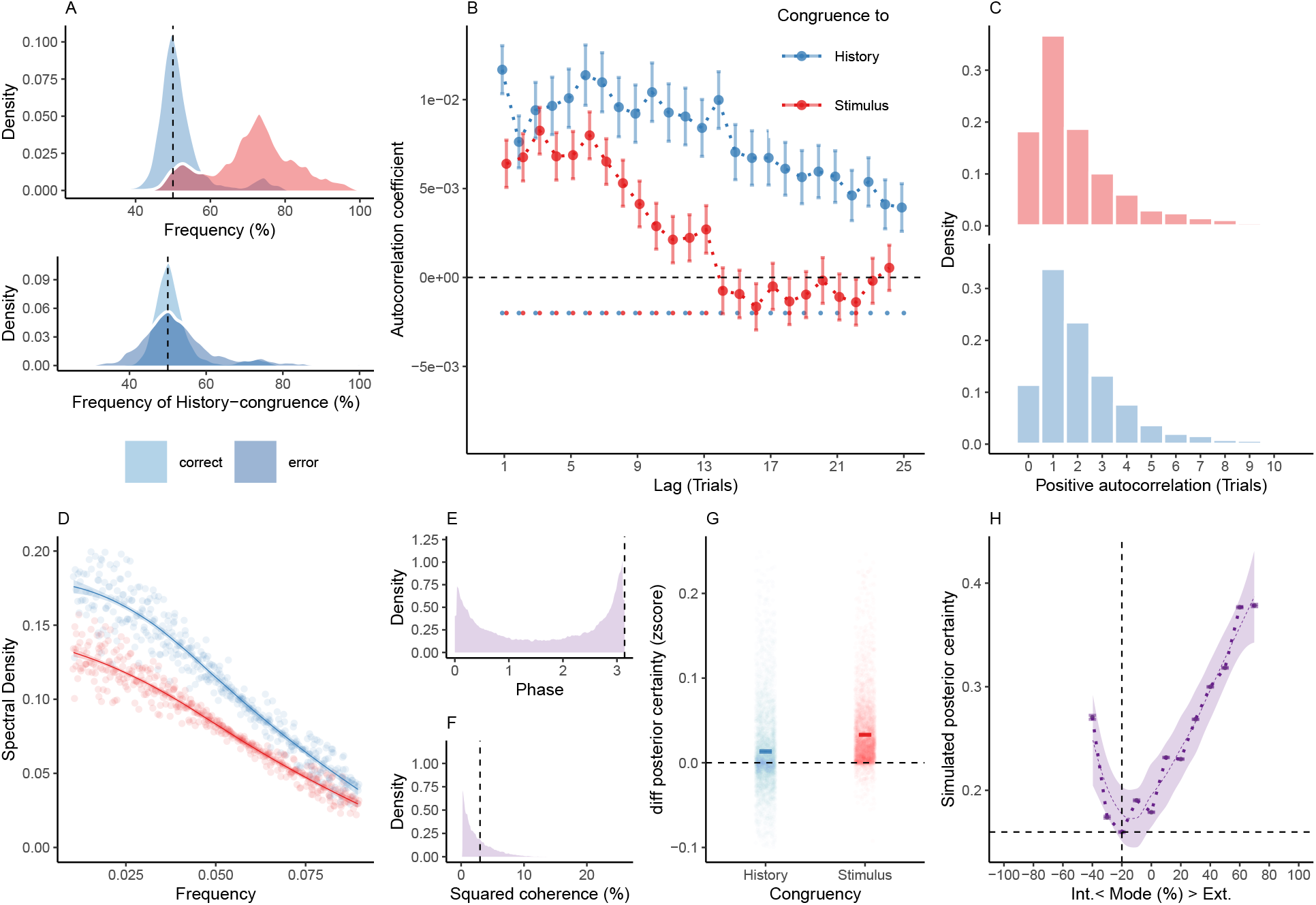
Internal and external modes in simulated perceptual decision-making. A. Simulated perceptual choices were stimulus-congruent in 71.36% ± 0.17% (in red) and history-congruent in 51.99% ± 0.11% of trials (in blue; T(4.32 × 10^3^) = 17.42, p = 9.89 × 10^-66^; upper panel). Due to the competition between stimulus- and history-congruence, history-congruent perceptual choices were more frequent when perception was stimulus-incongruent (i.e., on *error* trials; T(4.32 × 10^3^) = 11.19, p = 1.17 × 10^-28^; lower panel) and thus impaired performance in the randomized psychophysical design simulated here. B. At the simulated group level, we found significant autocorrelations in both stimulus-congruence (13 consecutive trials) and history-congruence (30 consecutive trials). C. On the level of individual simulated participants, autocorrelation coefficients exceeded the autocorrelation coefficients of randomly permuted data within a lag of 2.46 ± 1.17 × 10^-3^ trials for stimulus-congruence and 4.24 ± 1.85 × 10^-3^ trials for history-congruence. D. The smoothed probabilities of stimulus- and history-congruence (sliding windows of ±5 trials) fluctuated as *1/f noise*, i.e., at power densities that were inversely proportional to the frequency (power ∼ 1/*f ^β^*; stimulus-congruence: *β* = −0.81 ± 1.18 × 10^-3^, T(1.92 × 10^5^) = −687.58, p < 2.2 × 10^-308^; history-congruence: *β* = −0.83 ± 1.27 × 10^-3^, T(1.92 × 10^5^) = −652.11, p < 2.2 × 10^-308^). E. The distribution of phase shift between fluctuations in simulated stimulus- and history-congruence peaked at half a cycle (*π* denoted by dotted line). The dynamic probabilities of simulated stimulus- and history-congruence were therefore were strongly anti-correlated (*β* = −0.03 ± 8.22 × 10^-4^, T(2.12 × 10^6^) = −40.52, p < 2.2 × 10^-308^). F. The average squared coherence between fluctuations in simulated stimulus- and history-congruence (black dotted line) amounted to 6.49 ± 2.07 × 10^-3^ %. G. Simulated confidence was enhanced for stimulus-congruence (*β* = 0.03 ± 1.71 × 10^-4^, T(2.03 × 10^6^) = 178.39, p < 2.2 × 10^-308^) and history-congruence (*β* = 0.01 ± 1.5 × 10^-4^, T(2.03 × 10^6^) = 74.18, p < 2.2 × 10^-308^). H. In analogy to humans, the simulated data showed a quadratic relationship between the mode of perceptual processing and posterior certainty, which increased for stronger external and internal biases (*β*2 = 31.03 ± 0.15, T(2.04 × 10^6^) = 205.95, p < 2.2 × 10^-308^). The horizontal and vertical dotted lines indicate minimum posterior certainty and the associated mode, respectively.

To further probe the validity of the bimodal inference model, we tested whether posterior model quantities could explain aspects of the behavioral data that the model was not fitted to. First, we predicted that the posterior decision variable *Lt* not only encodes perceptual choices (i.e., the variable used for model estimation), but should also predict the speed of response and subjective confidence^30, 50^. Indeed, the estimated trial-wise posterior decision certainty |*L_t_*| correlated negatively with RTs in humans (*β* = −4.36 × 10^-3^ ± 4.64× 10^-4^, T(1.98 × 10^6^) = −9.41, p = 5.19 × 10^-21^) and TDs mice (*β* = −35.45 ± 0.86, T(1.28 × 10^6^) = −41.13, p < 2.2 × 10^-308^). Likewise, subjective confidence was positively correlated with the estimated posterior decision certainty in humans (*β* = 7.63 × 10^-3^ ± 8.32 × 10^-4^, T(2.06 × 10^6^) = 9.18, p = 4.48 × 10^-20^).

Second, the dynamic accumulation of information inherent to our model entails that biases toward perceptual history are stronger when the posterior decision certainty at the preceding trial is high^30, 31, 54^. Due to the link between posterior decision certainty and confidence, we reasoned that confident perceptual choices should be more likely to induce history-congruent perception at the subsequent trial^30, 31^. Indeed, logistic regression indicated that history-congruence was predicted by the posterior decision certainty |*Lt*-1 | (humans: *β* = 8.22 × 10^-3^ ± 1.94 × 10^-3^, z = 4.25, p = 2.17 × 10^-5^; mice: *β* = −3.72 × 10^-3^ ± 1.83 × 10^-3^, z = −2.03, p = 0.04) and subjective confidence (humans: *β* = 0.04 ± 1.62 × 10^-3^, z = 27.21, p = 4.56 × 10^-1^^63^) at the preceding trial.

In sum, computational modeling thus suggested that between-mode fluctuations are best explained by two interlinked processes (Figure 1E): (i), the dynamic accumulation of information across successive trials (i.e., following the estimated hazard rate *H*) and, (ii), ongoing anti-phase oscillations in the impact of external and internal information (i.e., as determined by *ωψ* and *ωLLR*).

### 5.8 Bimodal inference improves learning and perceptual metacognition in the absence of feedback

Is there a computational benefit to be gained from temporarily down-regulating biases toward preceding choices (Figure 2-3 B and C), instead of combining them with external sensory information at a constant weight (Supplemental Figure S7)? In their adaptive function for perceptual decision-making, internal predictions critically depend on error-driven learning to remain aligned with the current state of the world^56^. Yet when the same network processes external and internal information in parallel, inferences may become circular and maladaptive^57^: Ongoing decision-related activity may be distorted by noise in external sensory signals that are fed forward from the periphery or, alternatively, by aberrant internal predictions about the environment that are fed back form higher cortical levels^18, 57^.

Purely parallel processing therefore creates at least two challenges for perception: First, due to the sequential integration of inputs over time, internal predictions may progressively override sensory information^55^, leading to false inferences about the presented stimuli^19^. As a consequence, purely parallel processing may also lead to false inferences about the statistical regularities of volatile environments, where the underlying hazard rate *H = P(s_t_ = s_t_*_-1_) (i.e., the probability of a change in the state of the environment between two trials) may change over time. In the absence of feedback, agents have to update the estimate about *H* solely on the grounds of their experience, which is determined by the posterior log ratio *L_t_*. Yet *L_t_* depends not only on external information from the environment (the log likelihood ratio *LLRt*), but also on internal predictions, i.e., the log prior ratio *Lt*-1 and the assumed hazard rate *H_t_*. This circularity may impair the ability to learn about changes in *H* that occur in volatile environments (Figure 7A).

**Figure 7.**
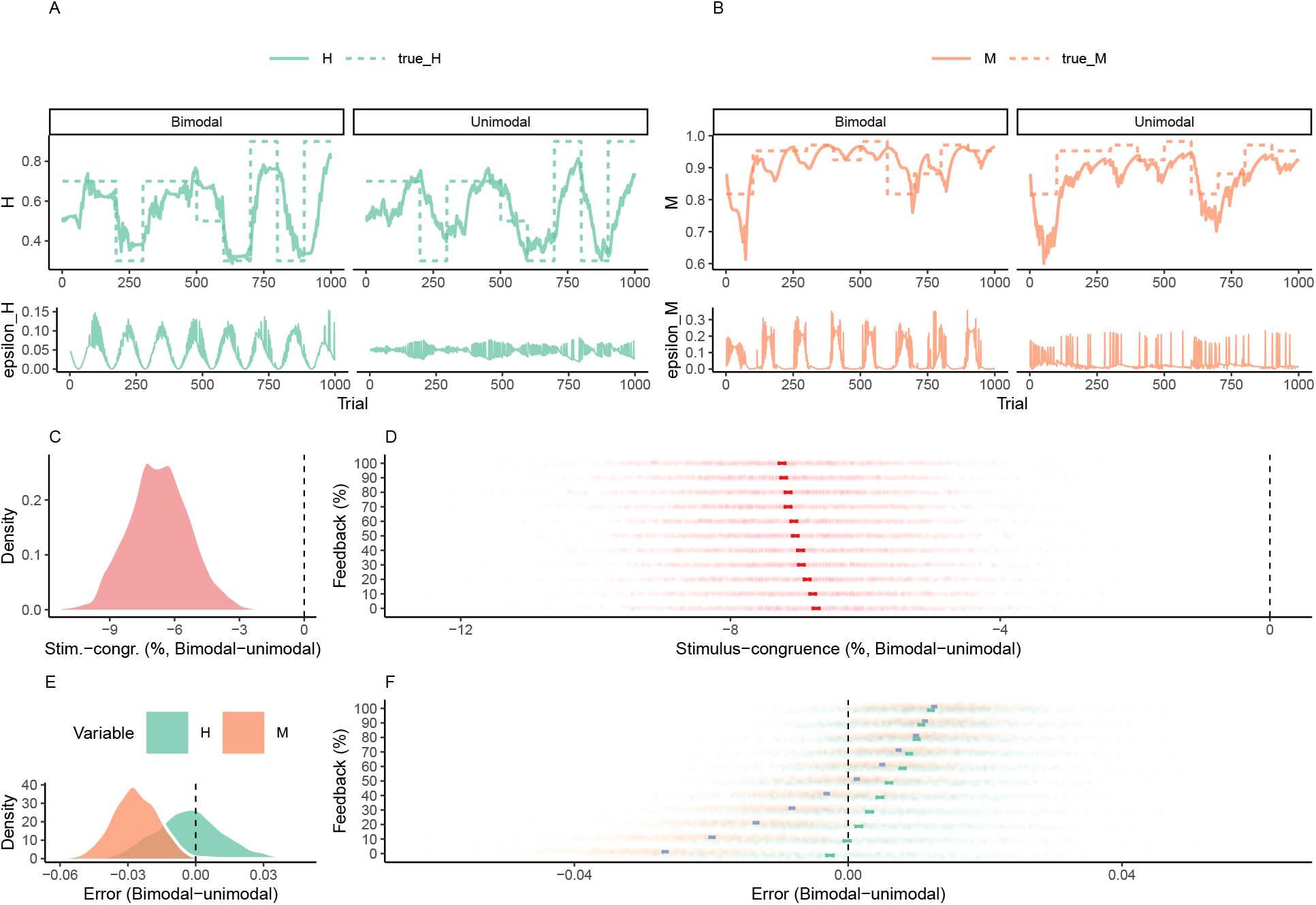
Adaptive benefits of bimodal inference. A. When the sensory environment changes unpredictably over time, agents have to update estimates *H_t_* (solid green line, upper panel) about the true hazard rate *H_t_* from experience (dotted green line, upper panel). Updates to *H_t_* are driven by an error term *e_H_* (solid green line, lower panel) that is defined by the difference between *Ht* and the presence of a perceived change in the environment. In contrast to the unimodal model (right panels), *e_H_* of the bimodal model (left panels) is modulated by a phasic component reflecting ongoing fluctuations between internal and external mode. B. When the precision of sensory encoding changes unpredictably over time, agents have to update estimates *Mt* (solid orange line, upper panel) about the true precision of sensory encoding *M_t_* from experience (dotted orange line, upper panel). Updates to *M_t_* are driven by an error term *e_M_* (red line, lower panel) that is defined by the difference between *M_t_* and the posterior decision-certainty. In contrast to the unimodal model (right panels), *< _M_* of the bimodal model (left panels) is modulated by a phasic component reflecting ongoing fluctuations between internal and external mode. C. In the absence of feedback, the bimodal inference model achieved lower stimulus-congruence as compared the unimodal control model (*β*1 = −6.71 ± 0.03, T(8.42 × 10^3^) = −234.31, p < 2.2 × 10^-308^). D. The unimodal control model benefited more strongly from the presence of external feedback, leading to a relative decrease in stimulus-congruence for the bimodal inference model at higher feedback levels (*β*2 = −0.05 ± 4.13 × 10^-3^, T(10 × 10^3^) = −12.32, p = 1.25 × 10^-34^). E. In the absence of feedback, the bimodal inference model achieved lower errors in the estimated hazard rate *H* (*β*_1_ = −2.94 × 10^-3^ ± 2.89 × 10^-4^, T(4.96 × 10^3^) = −10.18, p = 4.11× 10^-24^) as well as lower errors in the estimated probability of stimulus-congruent choices *M* (*β*1 = −0.03 ± 1.86 × 10^-4^, T(6.07 × 10^3^) = −137.75, p < 2.2 × 10^-308^). F. With an increasing availability of feedback, the advantage of the bimodal inference model was lost with respect to *H* (*β*_2_ = 1.43 × 10^-3^ ± 3.71 × 10^-5^, T(10 × 10^3^) = 38.58, p = 9.44×10^-304^)and*M* (*β*2 =3.91×10^-3^ ±2.51×10^-5^,T(10×10^3^)=156.18,p<2.2×10^-308^).

Second, purely parallel processing may also reduce the capacity to calibrate metacognitive beliefs about ongoing changes in the precision at which sensory signals are encoded. In the absence of feedback, agents depend on internal confidence signals^58^ (i.e., the absolute of the posterior log ratio |*Lt* |) to update beliefs *Mt* about the precision of sensory encoding *MI =* 1 — | *s_t_ — u_t_* |. While *Ml* depends only on the likelihood *LLR_t_,* the estimate *M_t_* is informed by the posterior *Lt*, which, in turn, is additionally modulated by the prior *Lt*-1 and *Ht*. Relying on internal predictions may thus distort metacognitive beliefs about the precision of sensory encoding (Figure 7B). This problem becomes particularly relevant when agents do not have full insight into the strength at which external and internal sources of information contribute to perceptual inference (i.e., when confidence is high during both internally- and externally-biased processing; Figure 2I-J; Figure 6G-H).

Here, we propose that bimodal inference may provide potential solutions to these problems of circular inference. By intermittently decoupling the decision variable *Lt* from internal predictions, between-mode fluctuations may create unambiguous error signals that adaptively update estimates about the hazard rate *Hl* and the precision of sensory encoding *M*. To illustrate this hypothesis, we simulated data for a total of 1000 participants who performed binary perceptual decisions for a total of 20 blocks of 100 trials each. Each block differed with respect to the true hazard rate *Hl* (either 0.1, 0.3, 0.5, 0.7 or 0.9) and the sensitivity parameter *α* (either 2, 3, 4, 5 or 6, determining *Ml* via the absolute of the log likelihood ratio | *LLRt*|, Figure 7A-B, upper panel). Importantly, the synthetic participants did not receive feedback on whether their perceptual decisions were correct.

We initialized each participant at a random value of *Hļ* (ranging from —0.25 to 0) and *Mļ* (ranging from 0.25 to 2), which were transformed into the unit interval to yield trial-wise estimates for *Ht* and *Mt*:

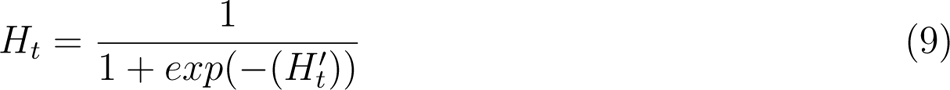

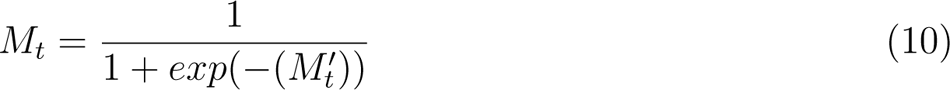

For each block, we generated stimuli *s_t_* using the true hazard rate *H.* Detected inputs *u_t_* were computed according to the block-wise sensitivity parameter *α*. Perceptual decisions *yt* were generated using the bimodal inference model with (*aψ* = *aLLR* = 1, *ζ* = 1 and *f* between 0.05 and 0.15) and a unimodal control model (*aψ* = *aLLR* = 0, *ζ* = 1).

Leaning about *H* was driven by the error-term *ε_H_* (Figure 7A, lower panel), reflecting the difference between *H_t_* and presence of a perceived change in the environment |*y_t_* − *y_t_*_-1_ |:

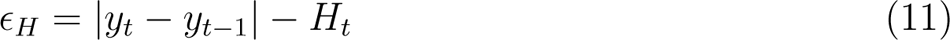

Trial-wise updates to *H* were provided by a Resorla-Wagner-rule with learning rate *βH* (ranging from 0.05 to 0.25). Since *y_t_* is more likely to accurately reflect the state of the environment during external mode, updates to *H* were additionally modulated by *ωLLR*:

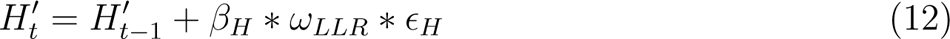

Learning about *M* was driven by error-term *ε_M_* (Figure 7B, lower panel), reflecting the difference between *Mt* and the posterior decision-certainty (1 *- \ yt − P* ( *yt* = 1) *\*):

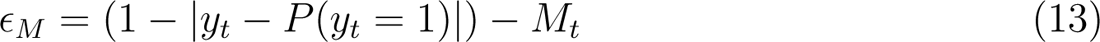

In analogy to *H*, we modeled trial-wise updates to *M* using a Rescorla-Wagner-rule with learning rate *β_M_* (ranging from 0.05 to 0.25). Since *y_t_* reflects the log likelihood ratio *LLR_t_* (and therefore the precision of sensory encoding) more closely during external mode, updates to *P* were additionally modulated by *ωLLR*:

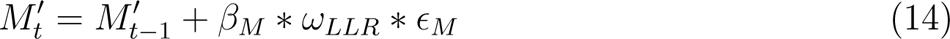

For each participant, we simulated data using both the bimodal inference model described above and a unimodal control model, in which the between-mode fluctuations were removed by setting the amplitude parameter *a* to zero (*aψ* = *aLLR* = 0). We compared the bimodal model of perceptual inference to the unimodal control model in terms of three dependent variables: the probability of stimulus-congruent perceptual choices, the error in the estimate about *H* (i.e., *H — H*|) and the error in the estimate about *M* (i.e., |*M* — *M*|, with *M =* 1 — (|*s_t_* — *u_t_*|)). We found that the bimodal inference model achieved lower stimulus-congruence in comparison to the unimodal control model (*B*_1_ = —6.71 ± 0.03, T(8.42×10^3^) = —234.31, p < 2.2×10^-308^, Figure 7C). At the same time, the bimodal inference model yielded lower errors in the estimated hazard rate *H* (*B*1 = —2.94×10^-3^ ± 2.89×10^-4^, T(4.96×10^3^)= —10.18, p =4.11×10^-24^) and probability of stimulus-congruent choices *P* (*B*1 = — 0.03 ± 1.86 × 10^-4^, T(6.07 × 10^3^) = — 137.75, p < 2.2 × 10^-308^, Figure 7E). This suggests that between-mode fluctuations may play an adaptive role for learning and perceptual metacognition by supporting robust inferences about the statistical regularities of volatile environments and ongoing changes in the precision of sensory encoding.

Finally, we asked whether differences between the bimodal inference model the unimodal control model depend on the presence of external feedback. We predicted that the benefits of the bimodal inference model over the unimodal control model should be lost when feedback is provided: With feedback, the error terms that induce updates in *H* and *P* can be informed by the true state of the environment *st* instead of posterior stimulus probabilities that are distorted by circular inferences:

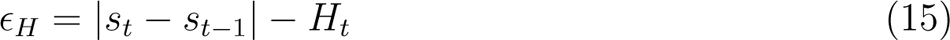

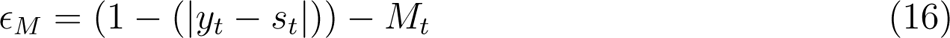

We repeated the above simulation for each participant while providing feedback on a subset of trials (10%, 20%, 30%, 40%, 50%, 60%, 70%, 80%, 90% and 100%). With increasing availability of external feedback, the bimodal inference model lost its advantage over the unimodal control model in terms of, (i), the estimated hazard rate *H* (*β*_2_ = 1.43 × 10^-3^ ± 3.71 × 10^-5^, T(10 × 10^3^) = 38.58, p = 9.44 × 10^-304^) and, (ii), the estimated probability of stimulus-congruent choices *M* (*β*_2_ = 3.91 × 10^-3^ ± 2.51 × 10^-5^, T(10 × 10^3^) = 156.18, p < 2.2 × 10^-308^, Figure 7F). This indicates that the benefits of bimodal inference are limited to situations in which external feedback is sparse.

## 6 Discussion

This work investigates the behavioral and computational characteristics of ongoing fluctuations in perceptual decision-making using two large-scale datasets in humans^20^ and mice^21^. We found that humans and mice cycle through recurring intervals of reduced sensitivity to external sensory information, during which they relied more strongly on perceptual history, i.e., an internal prediction that is provided by the sequence of preceding choices. Computational modeling indicated that these infra-slow periodicities are governed by two interlinked factors: (i), the dynamic integration of sensory inputs over time and, (ii), anti-phase oscillations in the strength at which perception is driven by internal versus external sources of information. These cross-species results suggest that ongoing fluctuations in perceptual decision-making arise not merely as a noise-related epiphenomenon of limited processing capacity, but result from a structured and adaptive mechanism that fluctuates between internally- and externally-oriented modes of sensory analysis.

### 6.1 Serial dependencies represent a pervasive and adaptive aspect of perceptual decision-making in humans and mice

A growing body of literature has highlighted that perception is modulated by preceding choices^22–28, 30, 32, 33^. Our work provides converging cross-species evidence supporting the notion that such serial dependencies are a pervasive and general phenomenon of perceptual decision-making (Figures 2 and 4, Supplemental Figures 1 and 3). While introducing errors in randomized psychophysical designs^24, 28, 30, 31, 43^ (Figures 2 and 4A), we found that perceptual history facilitates post-perceptual processes such as speed of response^42^ (Figure 2G) and subjective confidence in humans (Figure 2I).

At the level of individual traits, increased biases toward preceding choices were associated with reduced sensitivity to external information (Supplemental Figure 1C-D) and lower metacognitive efficiency. When investigating how serial dependencies evolve over time, we observed dynamic changes in strength of perceptual history (Figures 2 and 4B) that created wavering biases toward internally- and externally-biased modes of sensory processing. Between-mode fluctuations may thus provide a new explanation for ongoing changes in perceptual performance^6–11^.

In computational terms, serial dependencies may leverage the temporal autocorrelation of natural environments^31, 48^ to increase the efficiency of decision-making^35, 43^. Such temporal smoothing^48^ of sensory inputs may be achieved by updating dynamic predictions about the world based on the sequence of noisy perceptual experiences^22, 31^, using algorithms such as Kalman filtering^35^, Hierarchical Gaussian filtering^59^ or sequential Bayes^25, 42, 54^. At the level of neural mechanisms, the integration of internal with external information may be realized by combining feedback from higher levels in the cortical hierarchy with incoming sensory signals that are fed forward from lower levels^60^.

Yet relying too strongly on serial dependencies may come at a cost: When accumulating over time, internal predictions may eventually override external information, leading to circular and false inferences about the state of the environment. In this work, we used model simulations to show that, akin to the wake-sleep-algorithm in machine learning^61^, bimodal inference may help to determine whether errors result from external input or from internally-stored predictions (Figure 7): During internal mode, sensory processing is more strongly constrained by predictive processes that auto-encode the agent’s environment. Conversely, during external mode, the network is driven predominantly by sensory inputs^18^. Between-mode fluctuations may thus generate an unambiguous error signal that aligns internal predictions with the current state of the environment in iterative test-update-cycles^61^. On a broader scale, between-mode fluctuations may thus regulate the balance between feedforward versus feedback contributions to perception and thereby play a adaptive role in metacognition and reality monitoring^62^.

### 6.2 Arousal, attentional lapses, general response biases, insufficient training and metacognitive strategies as alternative explanations for between-mode fluctuations

These functional explanations for external and internal modes share the idea that, in order to form stable internal predictions about the statistical properties of the world (e.g., tracking the hazard rate of the environment) or metacognitive beliefs about processes occurring within the agent (e.g., monitoring ongoing changes in the reliability of feedback and feedforward processing), perception needs to temporarily disengage from internal predictions. By the same token, they presuppose that fluctuations in mode occur at the level of perceptual processing^26, 30, 48, 49^, and are not a passive phenomenon that is primarily driven by factors situated up- or downstream of sensory analysis.

First, it may be argued that agents stereotypically repeat preceding choices when less alert. Our analyses address this alternative driver of serial dependencies by building on the association between RTs and arousal^45, 47^. We found that RTs do not map linearly onto the mode of sensory processing, but become shorter for stronger biases toward both externally- and internally-oriented mode (Figure 2G-H; Figure 4I). These observations argue against the view that biases toward internal mode can be explained solely on the ground of ongoing changes in tonic arousal or fatigue^44^.

However, internal modes of sensory processing may also be attributed to attentional lapses^63^, which are caused by mind-wandering or mind-blanking and show a more complex relation to RTs^63^: While episodes of mind-blanking are characterized by an absence of subjective mental activity, more frequent misses, a relative increase in slow waves over posterior EEG electrodes and increased RTs, episodes of mind-wandering come along which rich inner experiences, more frequent false alarms, a relative increase of slow-wave amplitudes over frontal electrodes and decreased RTs^63^.

Yet in contrast to gradual between-mode fluctuations, engaging in mind-wandering as opposed to on-task attention seems to be an all-or-nothing phenomenon^63^. In addition, internally-biased processing did not increase either false alarms or misses, but induced choice errors through an enhanced impact of perceptual history (Figure 2 and 4A) that unfolded in alternating *streaks* ^9,13^ of elevated stimulus- and history-congruence. Finally, the increase in lapse rates during internal mode was not general, but history-dependent (Figures 3 and 5).

While these observations clearly distinguishes between-mode fluctuations from unspecific effects of lapses on decision-making, it remains an intriguing question for future research how mind-wandering and −blanking can be differentiated from internally-oriented modes of sensory processing in terms of their phenomenology, behavioral characteristics, neural signatures and noise profiles^10, 63^.

Second, it may be proposed that humans and mice apply a metacognitive response strategy that repeats preceding choices when less confident about their responses or when insufficiently trained on the task. In humans, however, confidence increased for stronger biases toward both external and internal mode (Figure 2I-J). For humans and mice, history-effects grew stronger with increasing exposure to (and expertise in) the task (Supplemental Figure S6). In addition, the existence of external and internal modes in murine perceptual decision-making (Figure 4) implies that between-mode fluctuations do not depend exclusively on the rich cognitive functions associated with human prefrontal cortex^64^.

Third, our computational modeling results provide further evidence against both of the above caveats: Simulations based on estimated model parameters closely matched the empirical data (Figure 6), reproduced aspects of behavior it was not fitted to (such as trial-wise confidence reports and RTs/TD for human and mice, respectively), and predicted that history-congruent choices occur more frequently after high-confidence trials^30, 31^. These findings suggest that perceptual choices and post-perceptual processes such as response behavior or metacognition are jointly driven by a dynamic decision variable^50^ that encodes uncertainty^31^ and is affected by ongoing changes in the integration of external versus internal information.

Of note, a recent computational study^65^ has used a Hidden Markov Model (HMM) to investigate perceptual decision-making in the IBL database^21^. In analogy to our findings, the authors observed that mice switch between temporally extended *strategies* that last for more than 100 trials: During *engaged* states, perception was highly sensitive to external sensory information. During *disengaged* states, in turn, choice behavior was prone to errors due to enhanced biases toward one of the two perceptual outcomes^65^. Despite the conceptual differences to our approach (discrete states in a HMM that correspond to switches between distinct decision-making strategies^65^ vs. gradual changes in mode that emerge from sequential Bayesian inference and ongoing fluctuations in the impact of external relative to internal information), it is tempting to speculate that engaged/disengaged states and between-mode fluctuations might tap into the same underlying phenomenon.

### 6.3 Fluctuations in mode as a driver of 1/f dynamics in perception

In light of the above, our results support the idea that, instead of unspecific effects of arousal, attention, training or metacognitive response strategies, perceptual choices are shaped by dynamic processes that occur at the level of sensory analysis^26, 30, 49^: (i), the integration of incoming signals over time and, (ii), ongoing fluctuations in the impact of external versus internal sources of decision-related information. It is particularly interesting that these two model components reproduce the established 1/f characteristic^36, 37^ of fluctuating performance in perception (see Figure 2-4D and previous work^9,10, 13^), since this feature has been attributed to both a memory process^13^ (corresponding to model component (i): internal predictions that are dynamically updated in response to new inputs) and wave-like variations in perceptual resources^9^ (corresponding to model component (ii): ongoing fluctuations in the impact of internal and external information).

1/f noise is an ubiquitous attribute of dynamic complex systems that integrate sequences of contingent sub-processes^36^ and exhibit self-organized criticality^37^. As most real-world processes are *critical*, i.e. not completely uniform (or subcritical) nor completely random (or supercritical)^37, 66^, the brain may have evolved to operate at a critical point as well^38^: Subcritical brains would be impervious to new inputs, whereas supercritical brains would be driven by noise. The 1/f observed in this study thus provides an intriguing connection between the notion that the brain’s self-organized criticality is crucial for balancing network stability with information transmission^38^ and the adaptive functions of between-mode fluctuations^18^, which we propose to support the build-up of robust internal predictions despite an ongoing stream of noisy sensory inputs.

### 6.4 Dopamine-dependent changes in E-I-balance as a neural mechanism of between-mode fluctuations

The link to self-organized criticality suggests that balanced cortical excitation and inhibition^67^ (E-I), which may enable efficient coding^67^ by maintaining neural networks in critical states^68^ could provide a potential neural mechanism of between-mode fluctuations. Previous work has proposed that the balance between glutamatergic excitation and GABA-ergic inhibition is regulated by activity-dependent feedback through NMDA receptors^69^. Such NMDA-mediated feedback has been related to the integration of external inputs over time^67^ (model component (i), Figure 1E), thereby generating serial dependencies in decision-making^70–73^. Intriguingly, slow neuromodulation by dopamine enhances NMDA-dependent signaling^70, 74, 75^ and fluctuates at infra-slow frequencies^76, 77^ that match the temporal dynamics of between-mode fluctuations observed in humans (Figure 2) and mice (Figure 4). Ongoing fluctuations in the impact of external versus internal information (model component (ii)) may thus by caused by phasic changes in E-I-balance that are induced by dopaminergic neuromodulation.

### 6.5 Limitations and open questions

In this study, we show that perception is attracted toward preceding choices in mice^21^ (Figure 4A) and humans (Figure 2A; see Supplemental Figure S1 for analyses within individual studies of the Confidence database^20^). Of note, previous work has shown that perceptual decision­making is concurrently affected by both attractive and repulsive serial biases that operate on distinct time-scales and serve complementary functions for sensory processing^27, 78, 79^: Short­term attraction may serve the decoding of noisy sensory inputs and increase the stability of perception, whereas long-term repulsion may enable efficient encoding and sensitivity to change^27^.

Importantly, repulsive biases operate in parallel to attractive biases^27^ and are therefore unlikely to account for the ongoing changes in mode that occur in alternating cycles of internally- and externally-oriented processing. To elucidate whether attraction and repulsion both fluctuate in their impact on perceptual decision-making will be an important task for future research, since this would help to understand whether attractive and repulsive biases are linked in terms of their computational function and neural implementation^27^.

A second open question concerns the neurobiological underpinnings of ongoing changes in mode. Albeit purely behavioral, our results tentatively suggest dopaminergic neuromodulation of NMDA-mediated feedback as one potential mechanism of externally- and internally-biased modes. Since between-mode fluctuations were found in both humans and mice, future studies can apply both non-invasive and invasive neuro-imaging and electrophysiology to better understand the neural mechanisms that generate ongoing changes in mode in terms of neuro-anatomy, −chemistry and −circuitry.

Finally, establishing the neural correlates of externally- an internally-biased modes will enable exiting opportunities to investigate their role for adaptive perception and decision­making. Causal interventions via pharmacological challenges, optogenetic manipulations or (non-)invasive brain stimulation will help to understand whether between-mode fluctuations are implicated in resolving credit-assignment problems^18, 80^ or in calibrating metacognition and reality monitoring^62^. Addressing these questions may therefore provide new insight into the pathophysiology of hallucinations and delusions, which have been characterized by an imbalance in the impact of external versus internal information^60, 81, 82^ and are typically associated with metacognitive failures and a departure from consensual reality^82^.

## 7 Methods

### 7.1 Ressource availability

#### 7.1.1 Lead contact

Further information and requests for resources should be directed to and will be fulfilled by the lead contact, Veith Weilnhammer.

#### 7.1.2 Materials availability

This study did not generate new unique reagents.

#### 7.1.3 Data and code availability

All custom code and behavioral data are available on https://github.com/veithweilnhammer/ Modes. This manuscript was created using the *R Markdown* framework, which integrates all data-related computations and the formatted text within one document. With this, we wish to make our approach fully transparent and reproducible for reviewers and future readers.

### 7.2 Experimental model and subject details

#### 7.2.1 Confidence database

We downloaded the human data from the Confidence database^20^ on 21/10/2020, limiting our analyses to the database category *perception*. Within this category, we selected studies in which participants made binary perceptual decision between two alternative outcomes (see Supplemental Table 1). We excluded two studies in which the average perceptual accuracy fell below 50%. After excluding these studies, our sample consisted of 21.05 million trials obtained from 4317 human participants and 66 individual studies.

#### 7.2.2 IBL database

We downloaded the murine data from the IBL database^21^ on 28/04/2021. We limited our analyses to the *basic task*, during which mice responded to gratings that appeared with equal probability in the left or right hemifield. Within each mouse, we excluded sessions in which perceptual accuracy was below 80% for stimuli presented at a contrast ≥ 50%. After exclusion, our sample consisted of 1.46 million trials trials obtained from N = 165 mice.

### 7.3 Method details

#### 7.3.1 Variables of interest

##### Primary variables of interest

We extracted trial-wise data on the presented stimulus and the associated perceptual decision. Stimulus-congruent choices were defined by perceptual decisions that matched the presented stimuli. History-congruent choices were defined by perceptual choices that matched the perceptual choice at the immediately preceding trial. The dynamic probabilities of stimulus- and history-congruence were computed in sliding windows of ±5 trials.

The *mode* of sensory processing was derived by subtracting the dynamic probability of history-congruence from the dynamic probability of stimulus-congruence, such that positive values indicate externally-oriented processing, whereas negative values indicate internally-oriented processing. When visualizing the relation of the mode of sensory processing to confidence, response times or trial duration (see below), we binned the mode variable in 10% intervals. We excluded bins than contained less than 0.5% of the total number of available data-points.

##### Secondary variables of interest

From the Confidence Database^20^, we furthermore extracted trial-wise confidence reports and response times (RTs; if RTs were available for both the perceptual decision and the confidence report, we only extracted the RT associated with the perceptual decision). To enable comparability between studies, we normalized RTs and confidence reports within individual studies using the *scale* R function. If not available for a particular study, RTs and confidence reports were treated as missing variables. From the IBL database^21^, we extracted trial durations (TDs) as defined by interval between stimulus onset and feedback, which represents a coarse measure of RT^21^.

##### Exclusion criteria for individual data-points

For non-normalized data (TDs from the IBL database^21^; d-prime, meta-dprime and M-ratio from the Confidence database^20^ and simulated confidence reports), we excluded data-points that differed from the median by more than 3 x MAD (median absolute distance^52^). For normalized data (RTs and confidence reports from the Confidence database^20^), we excluded data-points that differed from the mean by more than 3 x SD (standard deviation).

#### 7.3.2 Control variables

Next to the sequence of presented stimuli, we assessed the autocorrelation of task difficulty as an alternative explanation for any autocorrelation in stimulus- and history-congruence. For the Confidence Database^20^, task difficulty was indicated by one of the following labels: *Difficulty*, *Difference*, *Signal-to-Noise*, *Dot-Difference*, *Congruency*, *Coherence(-Level)*, *Dot-Proportion*, *Contrast(-Difference)*, *Validity*, *Setsize*, *Noise-Level(-Degree)* or *Temporal Distance*. When none of the above was available for a given study, task difficulty was treated as a missing variable. In analogy to RTs and confidence, difficulty levels were normalized within individual studies. For the IBL Database^21^, task difficulty was defined by the contrast of the presented grating.

#### 7.3.3 Autocorrelations

For each participant, trial-wise autocorrelation coefficients were estimated using the R-function *acf* with a maximum lag defined by the number of trials available per subject. Autocorrelation coefficients are displayed against the lag (in numbers of trials, ranging from 1 to 20) relative to the index trial (t = 0, see Figure 2B-C, 4B-C and 6B-C). To account for spurious autocorrelations that occur due to imbalances in the analyzed variables, we estimated autocorrelations for randomly permuted data (100 iterations). For group-level autocorrelations, we computed the differences between the true autocorrelation coefficients and the mean autocorrelation observed for randomly permuted data and averaged across participants.

At a given trial, group-level autocorrelation coefficients were considered significant when linear mixed effects modeling indicated that the difference between real and permuted autocorrelation coefficients was above zero at an alpha level of 0.05%. To test whether the autocorrelation of stimulus- and history-congruence remained significant when controlling for task difficulty and the sequence of presented stimuli, we added the respective autocorrelation as an additional factor to the linear mixed effects model that computed the group-level statistics (see also *Mixed effects modeling*).

To assess autocorrelations at the level of individual participants, we counted the number of subsequent trials (starting at the first trial after the index trial) for which less than 50% of the permuted autocorrelation coefficients exceeded the true autocorrelation coefficient. For example, a count of zero indicates that the true autocorrelation coefficients exceeded *less than 50%* of the autocorrelation coefficients computed for randomly permuted data at the first trial following the index trial. A count of five indicates that, for the first five trials following the index trial, the true autocorrelation coefficients exceeded *more than 50%* of the respective autocorrelation coefficients for the randomly permuted data; at the 6th trial following the index trial, however, *less than 50%* of the autocorrelation coefficients exceeded the respective permuted autocorrelation coefficients.

#### 7.3.4 Spectral analysis

We used the R function *spectrum* to compute the spectral densities for the dynamic probabilities of stimulus- and history-congruence as well as the phase (i.e., frequency-specific shift between the two time-series ranging from 0 to 2 * *π*) and squared coherence (frequency-specific variable that denotes the degree to which the shift between the two time-series in constant, ranging from 0 to 100%). Periodograms were smoothed using modified Daniell smoothers at a width of 50.

Since the dynamic probabilities of history- and stimulus-congruence were computed using a sliding windows of ±5 trials (i.e., intervals containing a total of 11 trials), we report the spectral density, coherence and phase for frequencies below 1/11 1/*Ntrials*. Spectral densities have one value per subject and frequency (data shown in Figures 2D anbd 4D). To assess the relation between stimulus- and history-congruence in this frequency range, we report average phase and average squared coherence for all frequencies below 1/11 1/*Ntrials* (i.e., one value per subject; data shown in Figure 2E-F and 4E-F).

Since the data extracted from the Confidence Database^20^ consist of a large set of individual studies that differ with respect to inter-trial intervals, we defined the variable *frequency* in the dimension of cycles per trial 1/*Ntrials* rather than cycles per second (Hz). For consistency, we chose 1/*Ntrials* as the unit of frequency for the IBL database^21^ as well.

### 7.4 Quantification and statistical procedures

All aggregate data are reported and displayed with errorbars as mean ± standard error of the mean.

#### 7.4.1 Mixed effects modeling

Unless indicated otherwise, we performed group-level inference using the R-packages *lmer* and *afex* for linear mixed effects modeling and *glmer* with a binomial link-function for logistic regression. We compared models based on Akaike Information Criteria (AIC). To account for variability between the studies available from the Confidence Database^20^, mixed modeling was conducted using random intercepts defined for each study. To account for variability across experimental session within the IBL database^21^, mixed modeling was conducted using random intercepts defined for each individual session. When multiple within-participant datapoints were analyzed, we estimated random intercepts for each participant that were *nested* within the respective study of the Confidence database^20^. By analogy, for the IBL database^21^, we estimated random intercepts for each session that were nested within the respective mouse. We report *β* values referring to the estimates provided by mixed effects modeling, followed by the respective T statistic (linear models) or z statistic (logistic models).

The effects of stimulus- and history-congruence on RTs and confidence reports (Figure 2, 4 and 6, subpanels G-I) were assessed in linear mixed effects models that tested for main effects of both stimulus- and history-congruence as well as the between-factor interaction. Thus, the significance of any effect of history-congruence on RTs and confidence reports was assessed while controlling for the respective effect of stimulus-congruence (and vice versa).

#### 7.4.2 Psychometric function

We obtained psychometric curves by fitting the following error function to the behavioral data:

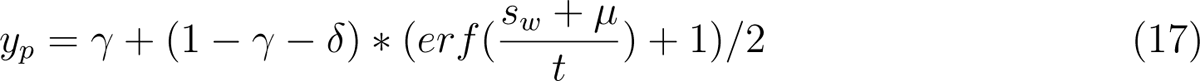

We used a maximum likelihood procedure to predict individual choices *y* (outcome A: *y* = 0; outcome B: *Y* = 1) from the choice probability *Yp*. In humans, we computed *sw* multiplying the inputs *s* (stimulus A: 0; outcome B: 1) with the task difficulty *Db* (binarized across 7 levels):

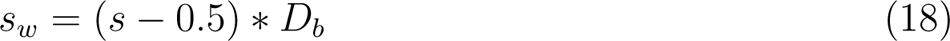

In mice, *s w* was defined by the respective stimulus contrast in the two hemifields:

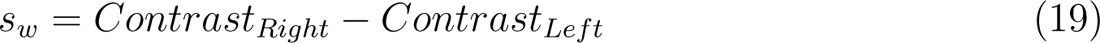

Parameters of the psychometric error function were fitted using the R-package *optimx*. The psychometric error function was defined via the parameters *γ* (lower lapse; lower bound = 0, upper bound = 0.5), *δ* (upper lapse; lower bound = 0, upper bound = 0.5), *µ* (bias; lower bound humans = −5; upper bound humans = 5, lower bound mice = −0.5, upper bound mice = 0.5) and threshold *t* (lower bound humans = 0.5, upper bound humans = 25; lower bound mice = 0.01, upper bound mice = 1.5).

#### 7.4.3 Computational modeling

##### Model definition

Our modeling analysis is an extension of a model proposed by Glaze et al.^54^, who defined a normative account of evidence accumulation for decision-making. In this model, trial-wise choices are explained by applying Bayes theorem to infer moment-by-moment changes in the state of environment from trial-wise noisy observations across trials.

Following Glaze et al.^54^, we applied Bayes rule to compute the posterior evidence for the two alternative choices (i.e., the log posterior ratio *L*) from the sensory evidence available at time-point *t* (i.e., the log likelihood ratio *LLR*) with the prior probability *ψ*:

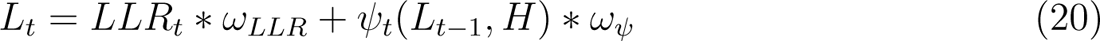

In the trial-wise design studied here, a transition between the two states of the environment (i.e., the sources generating the noisy observations available to the participant) can occur at any time. Despite the random nature of the psychophysical paradigms studied here^20, 21^, humans and mice showed significant biases toward preceding choices (Figure 2A and 4A). We thus assumed that the prior probability of the two possible outcomes depends on the posterior choice probability at the preceding trial and the hazard rate *H* assumed by the participant. Following Glaze et al.^54^, the prior *ψ* is thus computed as follows:

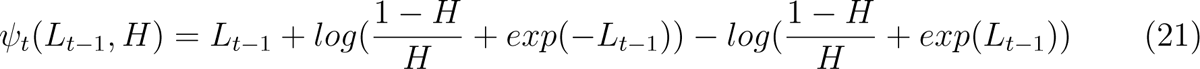

In this model, humans, mice and simulated agents make perceptual choices based on noisy observations *u*. The are computed by applying a sensitivity parameter *α* to the content of external sensory information *s*. For humans, we defined the input *s* by the two alternative states of the environment (stimulus A: *s* = 0; stimulus B: *s* = 1), which generated the observations *u* through a sigmoid function that applied a sensitivity parameter *α*:

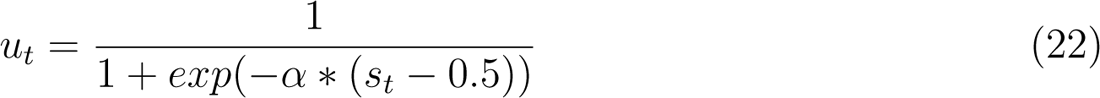

In mice, the inputs *s* were defined by the respective stimulus contrast in the two hemifields:

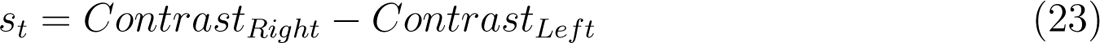

As in humans, we derived the input *u* by applying a sigmoid function with a sensitivity parameter *α* to input *s*:

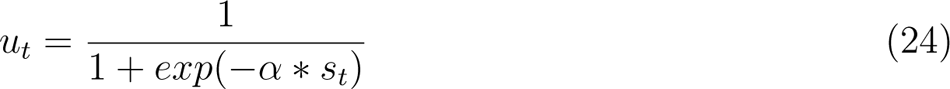

For humans, mice and in simulations, the log likelihood ratio *LLR* was computed from *u* as follows:

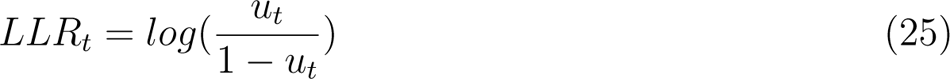

To allow for long-range autocorrelation in stimulus- and history-congruence (Figure 2B and 4B), our modeling approach differed from Glaze et al.^54^ in that it allowed for systematic fluctuation in the impact of sensory information (i.e., *LLR*) and the prior probability of choices *ψ* on the posterior probability *L*. This was achieved by multiplying the log likelihood ratio and the log prior ratio with coherent anti-phase fluctuations according to

##### Model fitting

In model fitting, we predicted the trial-wise choices *yt* (option A: 0; option B: 1) from inputs *s*. To this end, we minimized the log loss between *y_t_* and the choice probability *ypt* in the unit interval. *ypt* was derived from *Lt* using a sigmoid function defined by the inverse decision temperature *ζ*:

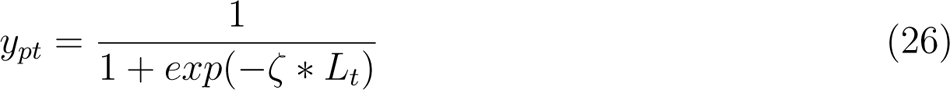

This allowed us to infer the free parameters *H* (lower bound = 0, upper bound = 1; human posterior = 0.45 ± 4.8 *×* 10^-5^; murine posterior = 0.46 ± 2.97 *×* 10^-4^), *α* (lower bound = 0, upper bound = 5; human posterior = 0.5 ± 1.12 *×* 10^-4^; murine posterior = 1.06 ± 2.88 *×* 10^-3^), *aψ* (lower bound = 0, upper bound = 10; human posterior = 1.44 ± 5.27 *×* 10^-4^; murine posterior = 1.71 ± 7.15 *×* 10^-3^), *ampLLR* (lower bound = 0, upper bound = 10; human posterior = 0.5 ± 2.02 *×* 10^-4^; murine posterior = 0.39 ± 1.08 *×* 10^-3^), frequency *f* (lower bound = 1/40, upper bound = 1/5; human posterior = 0.11 ± 1.68 *×* 10^-5^; murine posterior = 0.11 ± 1.63 *×* 10^-4^), *p* (lower bound = 0, upper bound = 2 *\* π*; human posterior = 2.72 ± 4.41 *×* 10^-4^; murine posterior = 2.83 ± 3.95 *×* 10^-3^) and inverse decision temperature *ζ* (lower bound = 1, upper bound = 10; human posterior = 4.63 ± 1.95 *×* 10^-4^; murine posterior = 4.82 ± 3.03 *×* 10^-3^) using the R-function *optimx*.

To validate our model, we correlated individual posterior parameter estimates with the respective conventional variables. We assumed that, (i), the estimated hazard rate *H* should correlate negatively with the frequency of history-congruent choices and that, (ii), the estimated *α* should correlate positively with the frequency of stimulus-congruent choices. In addition, we tested whether the posterior decision certainty (i..e. the absolute of the posterior log ratio) correlated negatively with RTs and positively with subjective confidence.

This allowed us to assess whether our model could explain aspects of the data it was not fitted to (i.e., RTs and confidence). Finally, we used simulations (see below) to show that all model components, including the anti-phase oscillations governed by *aψ*, *aLLR*, *f* and *p*, were necessary for our model to reproduce the empirical data observed for the Confidence database^20^ and IBL database^21^.

##### Model simulation 1: Data recovery

We used the posterior model parameters observed for humans (*H*, *α*, *aψ*, *aLLR* and *f*) to define individual parameters for simulation in 4317 simulated participants (i.e., equivalent to the number of human participants). For each participant, the number of simulated choices was drawn from a uniform distribution ranging from 300 to 700 trials. Inputs *s* were drawn at random for each trial, such that the sequence of inputs to the simulation did not contain any systematic seriality. Noisy observations *u* were generated by applying the posterior parameter *α* to inputs *s*, thus generating stimulus-congruent choices in 71.36 ± 2.6 × 10^-3^ % of trials. Choices were simulated based on the trial-wise choice probabilities *yp*. Simulated data were analyzed in analogy to the human and murine data. As a substitute of subjective confidence, we computed the absolute of the trial-wise posterior log ratio |*L*| (i.e., the posterior decision certainty).

##### Model simulation 2: Testing the adaptive benefits of bimodal inference

In contrast to the model applied to the behavioral data, our second set of simulations considered a situation in which agents learn about the properties of the environment from experience.

We modeled dynamic updates in the trial-wise estimates *Ht* about the true hazard rate *II = P(s_t_ = s_t_*_-1_) and trial-wise estimates *M_t_* about the precision of sensory encoding *M=\ −* (|*st* − *Utl)*.

In the absence of feedback, leaning about *Hì* was driven by the error-term *ε_H_*, which reflected the difference between the currently assumed hazard rate *H t* and the presence of a *perceived* change in the environment |*yt* − *yt*_ı|:

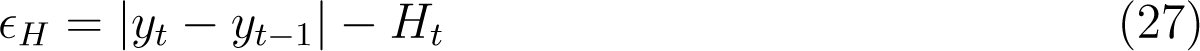

In the presence of feedback, *εH* reflected the difference between the currently assumed hazard rate *Ht* and an presence of a *true* change in the environment \ *st* − *st*-ı \:

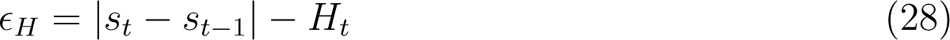

In the absence of feedback, learning about *M was* driven by the error-term *ε_M_,* reflecting the difference between *M t* and the posterior decision-certainty (1 − \ *yt* − *P* (*yt* = 1)\):

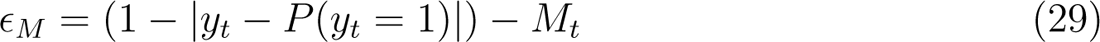

In the presence of feedback, *εM* reflected the difference between *M t* and the stimulus-congruence of the current response (1 − (\*yt* − *st*\)):

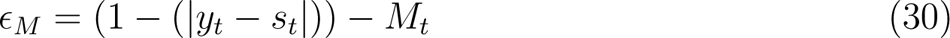

Updates to *H* and *M* were computed in logit-space using a Rescorla-Wagner-rule with learning rates defined by the product of *βH/M* and *ωLLR*. *Ht* and *M t* are computed by transforming *H* and *M’_t_* into the unit interval using a sigmoid function:

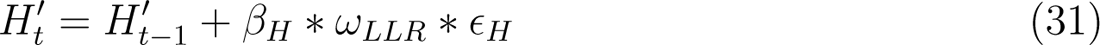

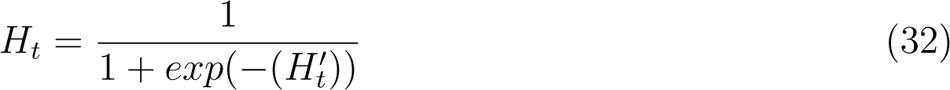

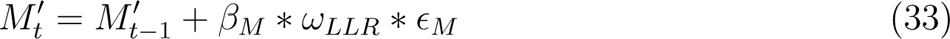

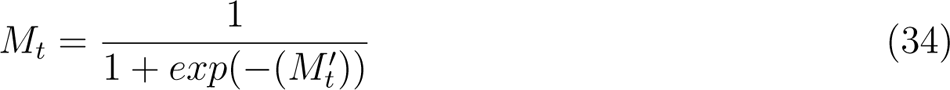

We simulated data for a total of 1000 participants for a total of 20 blocks of 100 trials each.

Each block differed with respect to the true hazard rate *H* (either 0.1, 0.3, 0.5, 0.7 or 0.9) and the sensitivity parameter *α* (either 2, 3, 4, 5 or 6, corresponding to values of *M* of 0.73, 0.82, 0.88, 0.92 or 0.95). Across participants, model parameters were set as follows: *H*_1_ initialized at random in a unit interval between −0.25 to 0; *P*_1_ initialized at random in a unit interval between 0.25 to 2; *a* = 1; *f* between 0.05 and 0.15 Hz; *ζ* = 1; *βH* and *βM* between 0.05 and 0.25. For each participant, we ran separate simulations with external feedback provided in 0%, 10%, 20%, 30%, 40%, 50%, 60%, 70%, 80%, 90% and 100% of trials.

## 9 Supplemental Items

**Supplemental Figure S1.**
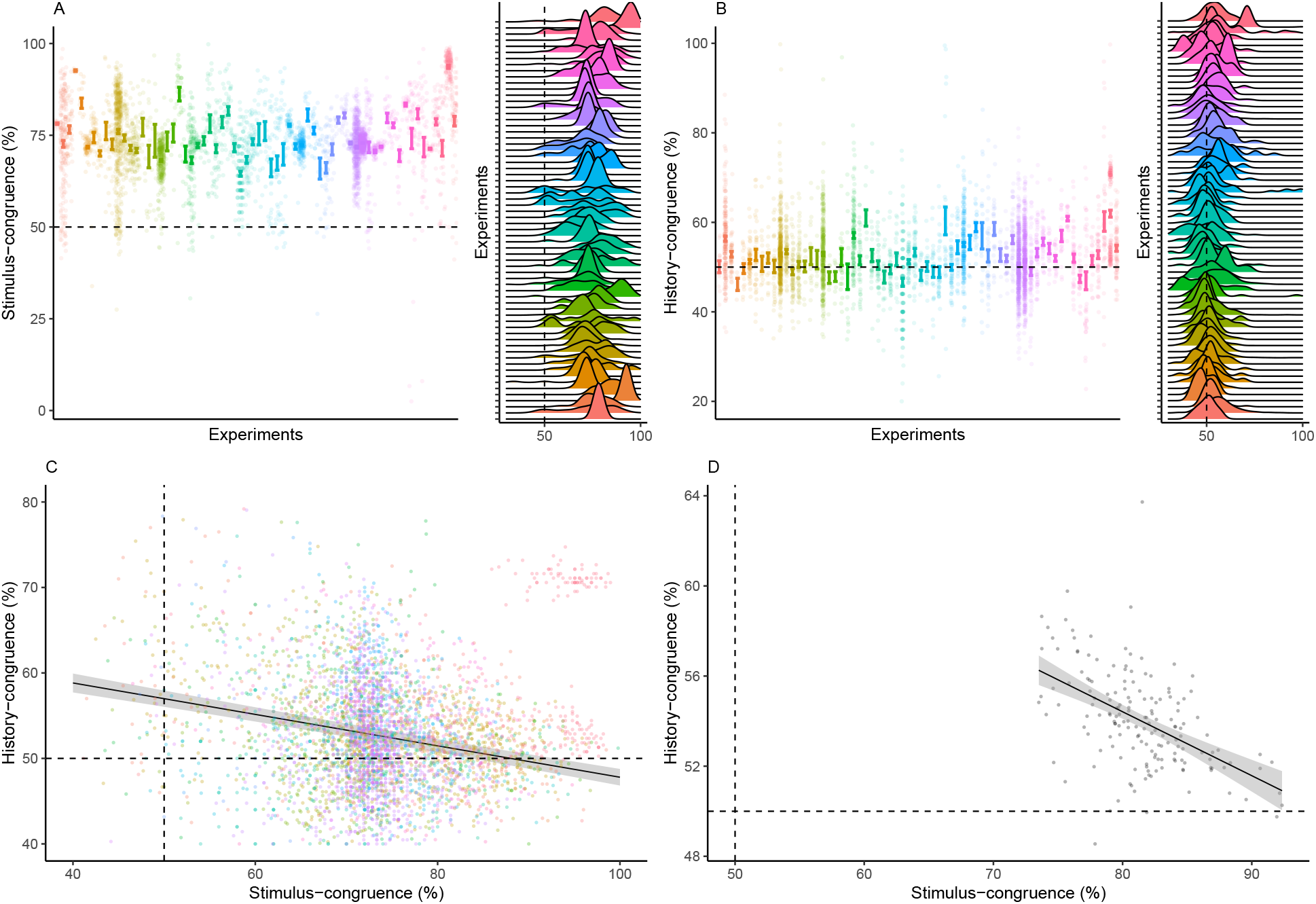
Stimulus- and history-congruence. A. Stimulus-congruent choices in humans amounted to 73.46% ± 0.15% of trials and were highly consistent across the experiments selected from the Confidence Database. B. History-congruent choices in humans amounted to 52.7% ± 0.12% of trials. In analogy to stimulus-congruence, the prevalence of history-congruence was highly consistent across the experiments selected from the Confidence Database. 48.48% of experiments showed significant (p < 0.05) attractive biases toward preceding choices, whereas 3.03% of experiments showed significant repulsive biases. C. In humans, we found an enhanced impact of perceptual history in participants who were less sensitive to external sensory information (T(4.3 × 10^3^) = −14.27, p = 3.78 × 10^-45^), suggesting that perception results from the competition of external with internal information. D. In analogy to humans, mice that were less sensitive to external sensory information showed stronger biases toward perceptual history (T(163) = −7.52, p = 3.44 × 10^-12^, Pearson correlation).

**Supplemental Figure S2.**
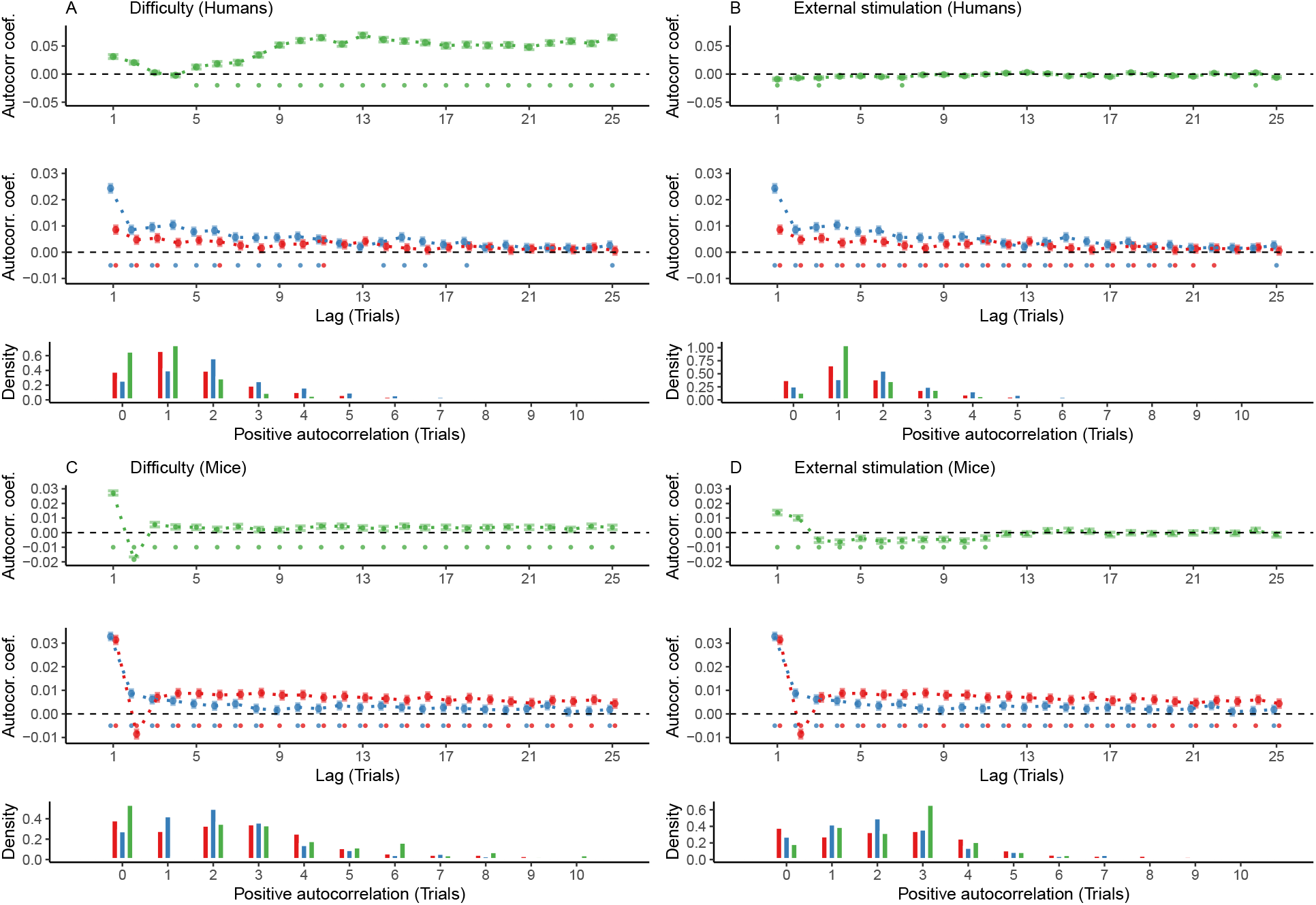
Controlling for task difficulty and external stimulation. In this study, we found highly significant autocorrelations of stimulus- and history-congruence in humans as well as in mice. Here, we show that these autocorrelations are not a trivial consequence of task difficulty or the sequence external stimulation. In addition, we computed trial-wise logistic regression coefficients as an alternative approach to assessing serial dependencies in stimulus- and history-congruence. A. In humans, task difficulty (in green) showed a significant autocorrelated starting at the 5th trial (upper panel, dots at the bottom indicate intercepts ≠ 0 in trial-wise linear mixed effects modeling at p < 0.05). When controlling for task difficulty, linear mixed effects modeling indicated a significant auto-correlation of stimulus-congruence (in red) for the first 3 consecutive trials (middle panel). 20% of trials within the displayed time window remained significantly autocorrelated. The autocorrelation of history-congruence (in blue) remained significant for the first 11 consecutive trials (64% significantly autocorrelated trials within the displayed time window). At the level of individual participants, the autocorrelation of task difficulty exceeded the respective autocorrelation of randomly permuted within a lag of 21.66 ± 8.37 × 10^-3^ trials (lower panel). B. The sequence of external stimulation (i.e., which of the two binary outcomes was supported by the presented stimuli; depicted in green) was negatively autocorrelated for 1 trial. When controlling for the autocorrelation of external stimulation, stimulus-congruence remained significantly autocorrelated for 22 consecutive trials (88% of trials within the displayed time window; lower panel) and history-congruence remained significantly autocorrelated for 20 consecutive trials (84% of trials within the displayed time window). At the level of individual participants, the autocorrelation of external stimulation exceeded the respective autocorrelation of randomly permuted within a lag of 2.94 ± 4.4 × 10^-3^ consecutive trials (lower panel). C. In mice, task difficulty showed an significant autocorrelated for the first 25 consecutive trials (upper panel). When controlling for task difficulty, linear mixed effects modeling indicated a significant auto-correlation of stimulus-congruence for the first 36 consecutive trials (middle panel). In total, 100% of trials within the displayed time window remained significantly autocorrelated. The autocorrelation of history-congruence remained significant for the first 8 consecutive trials, with 84% significantly autocorrelated trials within the displayed time window. At the level of individual mice, autocorrelation coefficients for difficulty were elevated above randomly permuted data within a lag of 15.13 ± 0.19 consecutive trials (lower panel). D. In mice, the sequence of external stimulation (i.e., which of the two binary outcomes was supported by the presented stimuli) was negatively autocorrelated for 11 consecutive trials (upper panel). When controlling for the autocorrelation of external stimulation, stimulus-congruence remained significantly autocorrelated for 86 consecutive trials (100% of trials within the displayed time window; middle) and history-congruence remained significantly autocorrelated for 8 consecutive trials (84% of trials within the displayed time window). At the level of individual mice, autocorrelation coefficients for external stimulation were elevated above randomly permuted data within a lag of 2.53 ± 9.8 × 10^-3^ consecutive trials (lower panel).

**Supplemental Figure S3.**
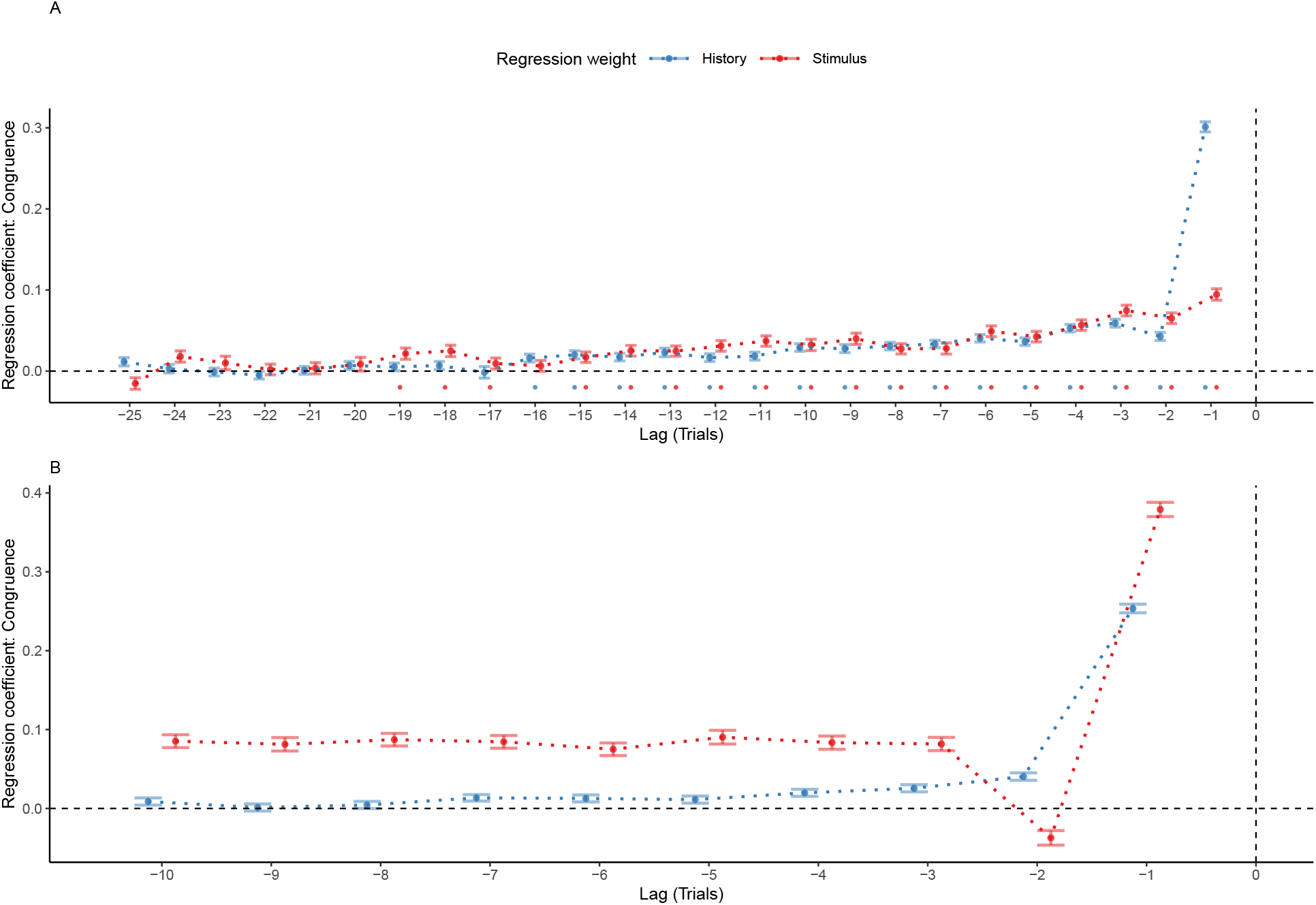
Reproducing group-level autocorrelations using logistic regression. A. As an alternative to group-level autocorrelation coefficients, we used trial-wise logistic regression to quantify serial dependencies in stimulus- and history-congruence. This analysis predicted stimulus- and history-congruence at the index trial (trial *t* = 0, vertical line) based on stimulus- and history-congruence at the 25 preceding trials. Mirroring the shape of the group-level autocorrelations, trial-wise regression coefficients (depicted as mean ± SEM, dots mark trials with regression weights significantly greater than zero at p < 0.05) increased toward the index trial *t* = 0 for the human data. B. Following our results in human data, regression coefficients that predicted history-congruence at the index trial (trial t = 0, vertical line) increased exponentially for trials closer to the index trial in mice. In contrast to history-congruence, stimulus-congruence showed a negative regression weight (or autocorrelation coefficient, see Figure 4B) at trial −2. This was due to the experimental design (see also the autocorrelations of difficulty and external stimulation in Supplemental Figure S2C and D): When mice made errors at easy trials (contrast ≥ 50%), the upcoming stimulus was shown at the same spatial location and at high contrast. This increased the probability of stimulus-congruent perceptual choices after stimulus-incongruent perceptual choices at easy trials, thereby creating a negative regression weight (or autocorrelation coefficient) of stimulus-congruence at trial −2.

**Supplemental Figure S4.**
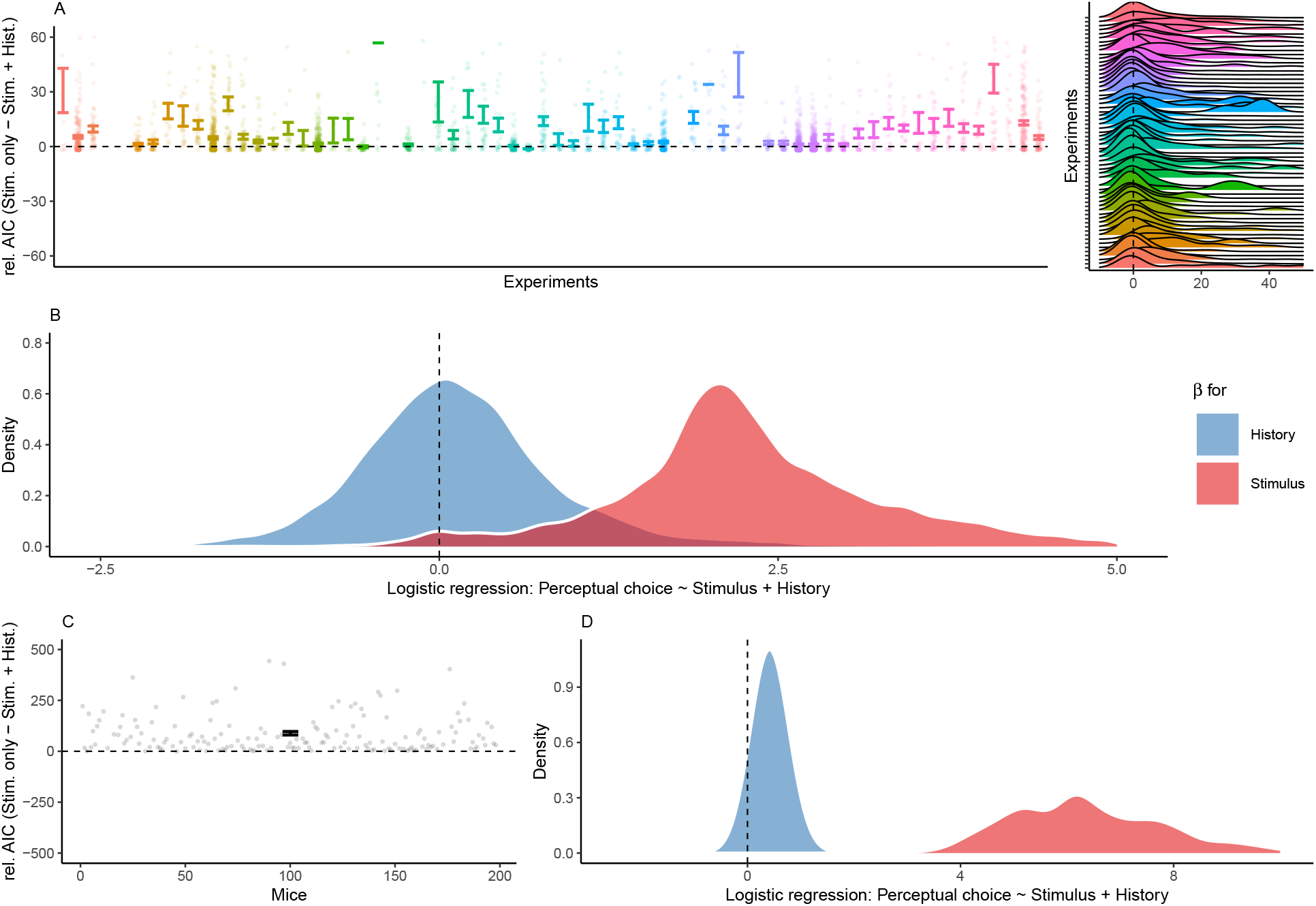
History-congruence in logistic regression. A. To ensure that perceptual history played a significant role in perception despite the ongoing stream of external information, we tested whether human perceptual decision-making was better explained by the combination of external and internal information or, alternatively, by external information alone. To this end, we compared Aikake information criteria between logistic regression models that predicted trial-wise perceptual responses either by both current external sensory information and the preceding percept, or by external sensory information alone (values above zero indicate a superiority of the full model). With high consistency across the experiments selected from the Confidence Database, this model-comparison confirmed that perceptual history contributed significantly to perception (difference in AIC = 8.07 ± 0.53, T(57.22) = 4.1, p = 1.31 × 10^-4^). B. Participant-wise regression coefficients amount to 0.18 ± 0.02 for the effect of perceptual history and 2.51 ± 0.03 for external sensory stimulation. C. In mice, an AIC-based model comparison indicated that perception was better explained by logistic regression models that predicted trial-wise perceptual responses based on both current external sensory information and the preceding percept (difference in AIC = 88.62 ± 8.57, T(164) = −10.34, p = 1.29 × 10^-19^). D. In mice, individual regression coefficients amounted to 0.42 ± 0.02 for the effect of perceptual history and 6.91 ± 0.21 for external sensory stimulation.

**Supplemental Figure S5.**
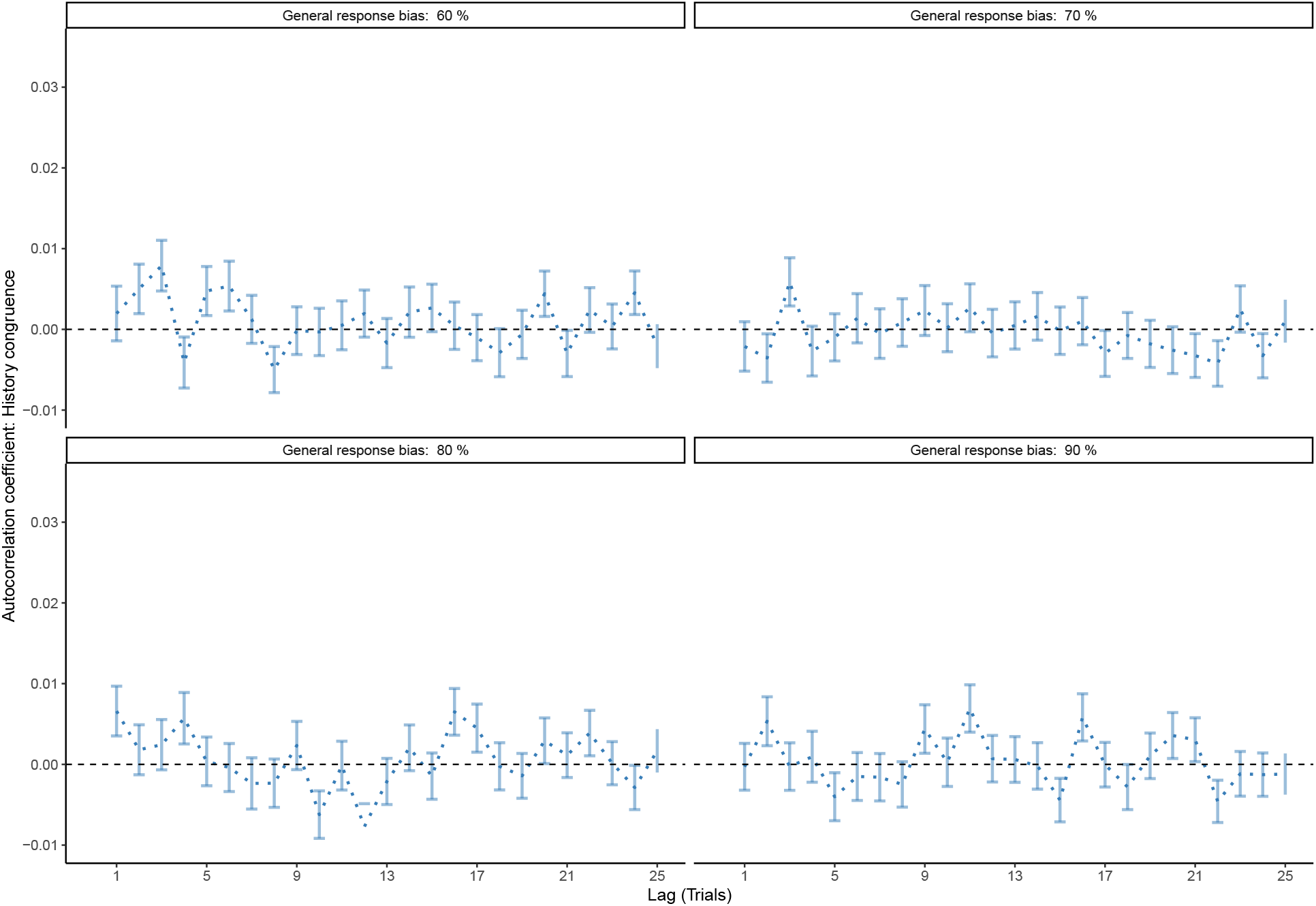
Correcting for general response biases. Here, we ask whether the autocorrelation of history-congruence (as shown in Figure 2-3C) may be driven by general response biases (i.e., a general propensity to choose one of the two possible outcomes more frequently than the alternative). To this end, we generated sequences of 100 perceptual choices with general response biases ranging from 60 to 90% for 1000 simulated participants each. We then computed the autocorrelation of history-congruence for these simulated data. Crucially, we used the correction procedure that is applied to all autocorrelation curves shown in this manuscript: All reported autocorrelation coefficients are computed relative to the average autocorrelation coefficients obtained for 100 iterations of randomly permuted trial sequences. The above simulation show that this correction procedure removes any potential contribution of general response biases to the auto-correlation of history-congruence. This indicates that the autocorrelation of history-congruence (as shown in Figure 2-3C) is not driven by general response biases that were present in the empirical data at a level of 58.71% ± 0.22% in humans and 54.6% ± 0.3% in mice.

**Supplemental Figure S6.**
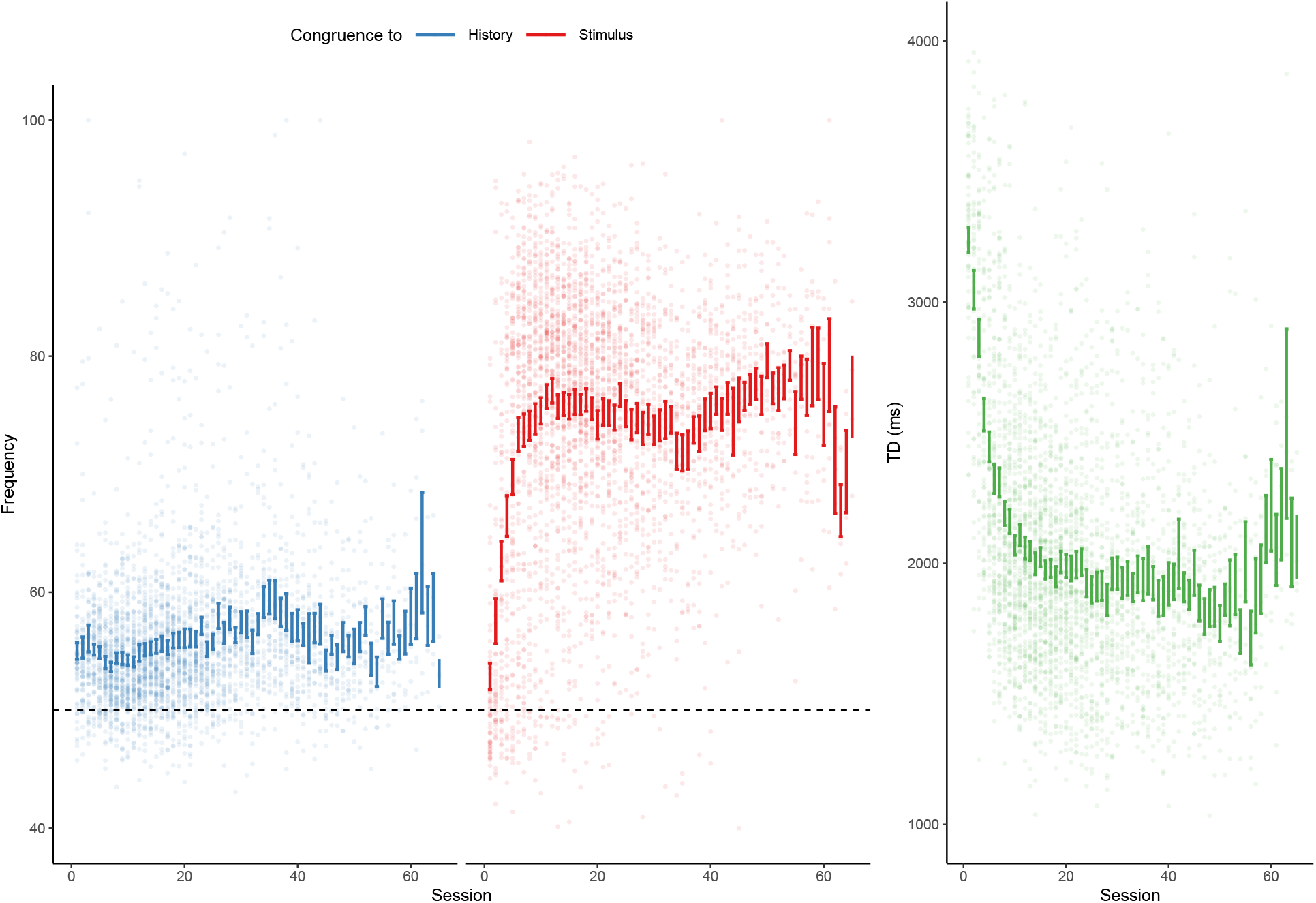
History-/stimulus-congruence and TDs during training of the basic task. Here, we depict the progression of history- and stimulus-congruence (depicted in blue and red, respectively; left panel) as well as TDs (in green; right panel) across training sessions in mice that achieved proficiency (i.e., stimulus-congruence ≥ 80%) in the *basic* task of the IBL dataset. We found that both history-congruent perceptual choices (*β* = 0.13 ± 4.67 × 10^-3^, T(8.4 × 10^3^) = 27.04, p = 1.96 × 10^-154^) and stimulus-congruent perceptual choices (*β* = 0.34 ± 7.13 × 10^-3^, T(8.51 × 10^3^) = 47.66, p < 2.2 × 10^-308^) became more frequent with training. As in humans, mice showed shorter TDs with increase exposure to the task (*β* = −22.14 ± 17.06, T(1.14 × 10^3^) = −1.3, p < 2.2 × 10^-308^).

**Supplemental Figure S7.**
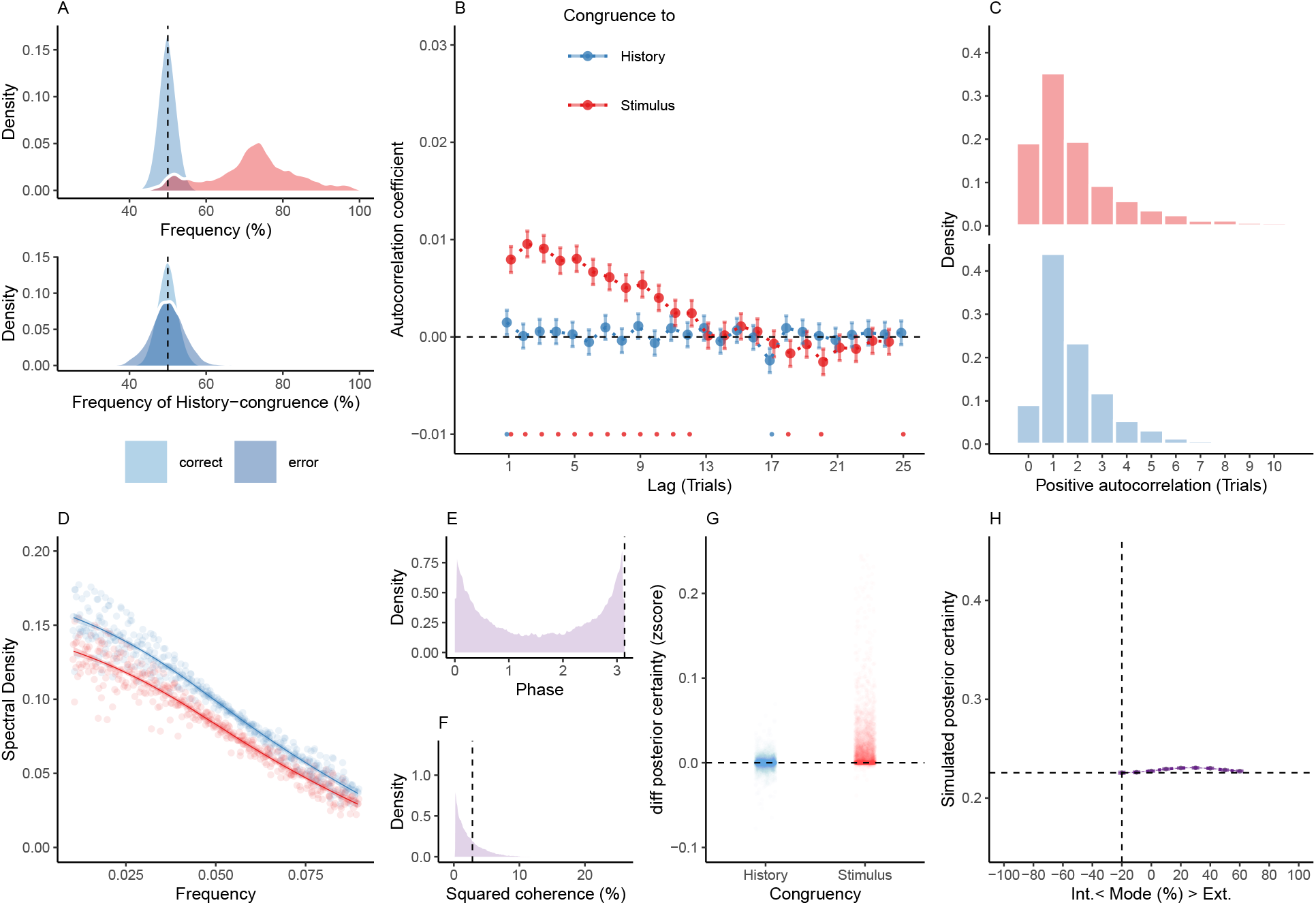
Reduced Control Model 1: No accumulation of information across trials. When simulating data for the *no-accumulation model*, we removed the accumulation of information across trials by setting the Hazard rate *H* to 0.5. Simulated data thus depended only on the participant-wise estimates for the amplitudes *aLLR/ψ*, frequency *f*, phase *p* and inverse decision temperature *ζ*. A. Similar to the full model (Figure 6), simulated perceptual choices were stimulus-congruent in 72.14% ± 0.17% of trials (in red). History-congruent amounted to 49.89% ± 0.03% of trials (in blue). In contrast to the full model, the no-accumulation model showed a significant bias against perceptual history T(4.32 × 10^3^) = −3.28, p = 1.06 × 10^-3^; upper panel). In contrast to the full model, there was no difference in the frequency of history-congruent choices between correct and error trials (T(4.31 × 10^3^) = 0.76, p = 0.44; lower panel). B. In the no-accumulation model, we found no significant autocorrelation of history-congruence beyond the first trial, whereas the autocorrelation of stimulus-congruence was preserved. C. In the no-accumulation model, the number of consecutive trials at which true autocor­relation coefficients exceeded the autocorrelation coefficients for randomly permuted data increased with respect to stimulus-congruence (2.83 ± 1.49 × 10^-3^ trials; T(4.31 × 10^3^) = 3.45, p = 5.73 × 10^-4^) and decreased with respect to history-congruence (1.85 ± 3.49 × 10^-4^ trials; T(4.32 × 10^3^) = −19.37, p = 3.49 × 10^-80^) relative to the full model. D. In the no-accumulation model, the smoothed probabilities of stimulus- and history-congruence (sliding windows of ±5 trials) fluctuated as *1/f noise*, i.e., at power densities that were inversely proportional to the frequency (power ∼ 1/*f ^β^*; stimulus-congruence: *β* = −0.82 ± 1.2 × 10^-3^, T(1.92 × 10^5^) = −681.98, p < 2.2 × 10^-308^; history-congruence: *β* = −0.78 ± 1.11 × 10^-3^, T(1.92 × 10^5^) = −706.57, p < 2.2 × 10^-308^). E. In the no-accumulation model, the distribution of phase shift between fluctuations in simulated stimulus- and history-congruence peaked at half a cycle (*π* denoted by dotted line). In contrast to the full model, the dynamic probabilities of simulated stimulus- and history-congruence were not significantly anti-correlated (*β* = 6.39 × 10^-4^ ± 7.22 × 10^-4^, T(8.89 × 10^5^) = 0.89, p = 0.38). F. In the no-accumulation model, the average squared coherence between fluctuations in simulated stimulus- and history-congruence (black dotted line) was reduced in comparison to the full model (T(3.56 × 10^3^) = −9.96, p = 4.63 ×10^-23^) and amounted to 2.8 ± 7.29 × 10^-4^%. G. Similar to the full model, confidence simulated from the no-accumulation model was enhanced for stimulus-congruent choices (*β* = 0.01 ± 9.4 × 10^-5^, T(2.11 × 10^6^) = 158.1, p < 1. 2.2 × 10^-308^). In contrast to the full model (Figure 6), history-congruent choices were not characterized by enhanced confidence (*β* = 8.78 × 10^-5^ ± 8.21 × 10^-5^, T(2.11 × 10^6^) = 1.07, p = 0. 29). H. In the no-accumulation model, the positive quadratic relationship between the mode of perceptual processing and confidence was markedly reduced in comparison to the full model (*β*2 = 0.19 ± 0.06, T(2.11 × 10^6^) = 3, p = 2.69 × 10^-3^). The horizontal and vertical dotted lines indicate minimum posterior certainty and the associated mode, respectively.

**Supplemental Figure S8.**
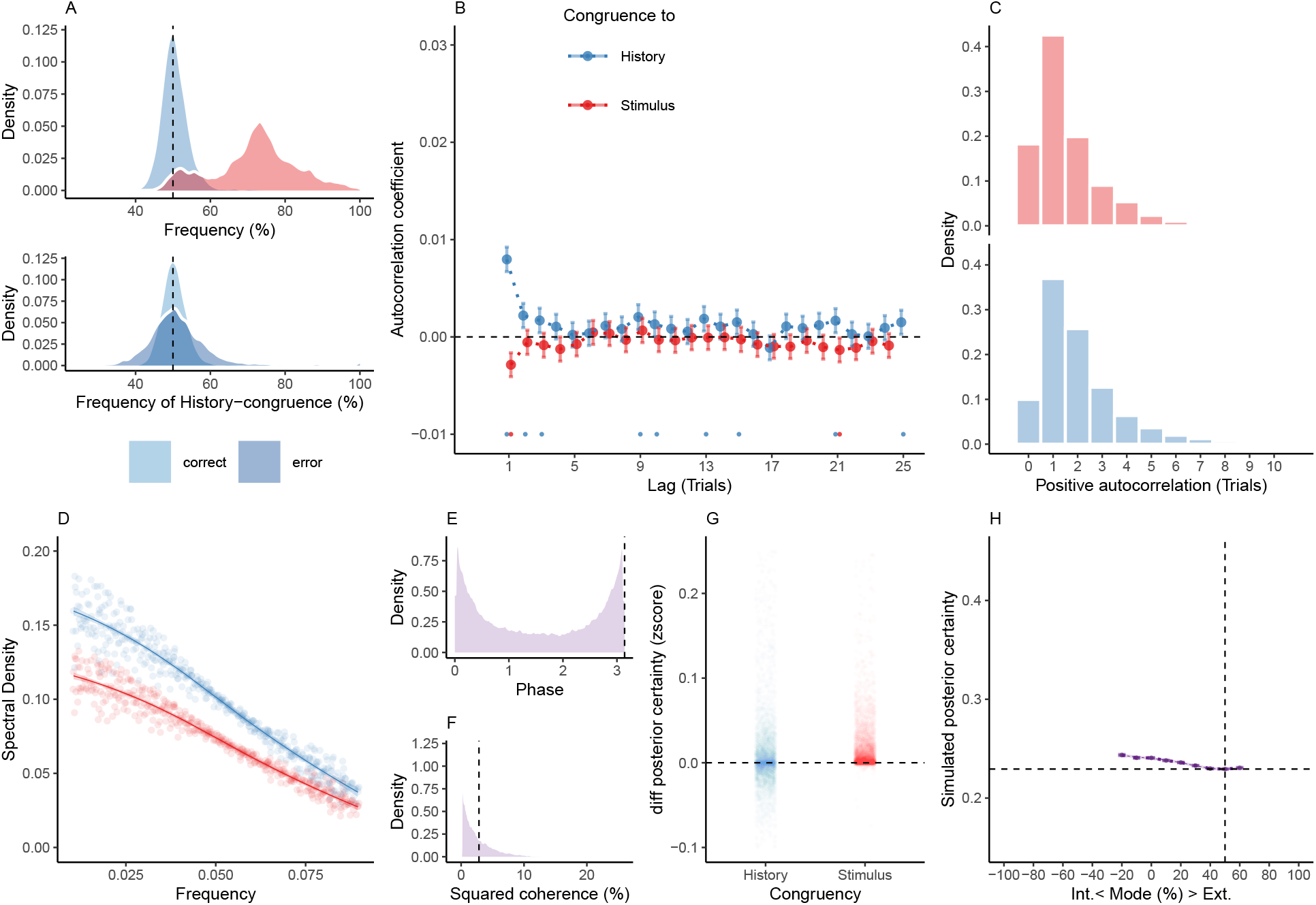
Reduced Control Model 2: No oscillations. When simulating data for the *no-oscillation model*, we removed the oscillation from the likelihood and prior terms by setting the amplitudes *aLLR* and *aψ* to zero. Simulated data thus depended only on the participant-wise estimates for hazard rate *H* and inverse decision temperature *ζ*. A. Similar to the full model (Figure 6), simulated perceptual choices were stimulus-congruent in 71.97% ± 0.17% of trials (in red). History-congruent amounted to 50.73% ± 0.07% of trials (in blue). As in the full model, the no-oscillation model showed a significant bias toward perceptual history T(4.32 × 10^3^) = 9.94, p = 4.88 × 10^-23^; upper panel). Similarly, history-congruent choices were more frequent at error trials (T(4.31 × 10^3^) = 10.59, p = 7.02 × 10^-26^; lower panel). B. In the no-oscillation model, we did not find significant autocorrelations for stimulus-congruence. Likewise, we did not observe any autocorrelation of history-congruence beyond the first three consecutive trials. C. In the no-oscillation model, the number of consecutive trials at which true autocorrelation coefficients exceeded the autocorrelation coefficients for randomly permuted data decreased with respect to both stimulus-congruence (1.8 ± 1.59 × 10^-3^ trials; T(4.31 × 10^3^) = −5.21, p = 2 × 10^-7^) and history-congruence (2.18 ± 5.48 × 10^-4^ trials; T(4.32 × 10^3^) = −17.1, p = 75 × 10^-63^) relative to the full model. D. In the no-oscillation model, the smoothed probabilities of stimulus- and history-congruence (sliding windows of ±5 trials) fluctuated as *1/f noise*, i.e., at power densities that were inversely proportional to the frequency (power ∼ 1/*f ^β^*; stimulus-congruence: *β* = −0.78 ± 1.1 × 10^-3^, T(1.92 × 10^5^) = −706.93, p < 2.2 × 10^-308^; history-congruence: *β* = −0.79 ± 1.12 × 10^-3^, T(1.92 × 10^5^) = −702.46, p < 2.2 × 10^-308^). E. In the no-oscillation model, the distribution of phase shift between fluctuations in simulated stimulus- and history-congruence peaked at half a cycle (*π* denoted by dotted line). In contrast to the full model, the dynamic probabilities of simulated stimulus- and history-congruence were positively correlated (*β* = 4.3× 10^-3^ ± 7.97× 10^-4^, T(1.98 ×10^6^) = 5.4, p = 6.59× 10^-8^). F. In the no-oscillation model, the average squared coherence between fluctuations in simulated stimulus- and history-congruence (black dottet line) was reduced in comparison to the full model (T(3.52 × 10^3^) = −6.27, p = 3.97 × 10^-10^) and amounted to 3.26 ± 8.88 × 10^-4^%. G. Similar to the full model, confidence simulated from the no-oscillation model was enhanced for stimulus-congruent choices (*β* = 0.01 ± 1.05× 10^-4^, T(2.1 ×10^6^) = 139.17, p < 2.2×10^-308^) and history-congruent choices (*β* = 8.05 × 10^-3^ ± 9.2 × 10^-5^, T(2.1 × 10^6^) = 87.54, p < 2.2 × 10^-308^). H. In the no-oscillation model, the positive quadratic relationship between the mode of perceptual processing and confidence was markedly reduced in comparison to the full model (*β*2 = 0.14 ± 0.07, T(2.1 × 10^6^) = 1.95, p = 0.05). The horizontal and vertical dotted lines indicate minimum posterior certainty and the associated mode, respectively.

**Supplemental Figure S9.**
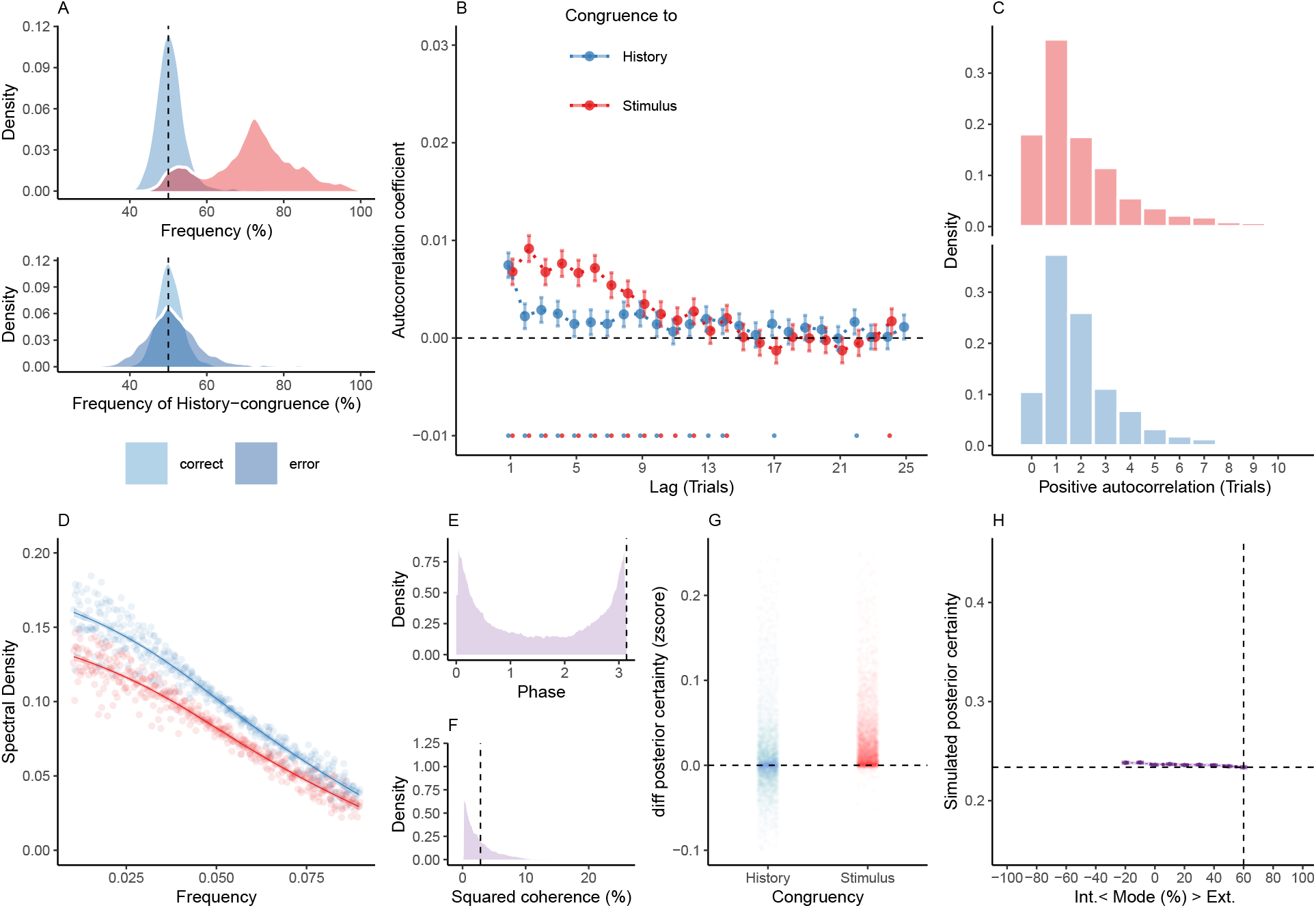
Reduced Control Model 3: Only oscillation of the likelihood. When simulating data for the *likelihood-oscillation-only model*, we removed the oscillation from the prior term by setting the amplitude *aψ* to zero. Simulated data thus depended only on the participant-wise estimates for hazard rate *H*, amplitude *aLLR*, frequency *f*, phase *p* and inverse decision temperature *ζ*. A. Similar to the full model (Figure 6), simulated perceptual choices were stimulus-congruent in 71.97% ± 0.17% of trials (in red). History-congruent amounted to 50.76% ± 0.07% of trials (in blue). As in the full model, the likelihood-oscillation-only model showed a significant bias toward perceptual history T(4.32 × 10^3^) = 10.29, p = 1.54 × 10^-24^; upper panel). Similarly, history-congruent choices were more frequent at error trials (T(4.32 × 10^3^) = 9.71, p = 4.6 × 10^-22^; lower panel). B. In the likelihood-oscillation-only model, we observed that the autocorrelation coefficients for history-congruence were reduced below the autocorrelation coefficients of stimulus-congruence. This is an approximately five-fold reduction relative to the empirical results observed in humans (Figure 2B), where the autocorrelation of history-congruence was above the autocorrelation of stimulus-congruence. Moreover, in the reduced model shown here, the number of consecutive trials that showed significant autocorrelation of history-congruence was reduced to 11. C. In the likelihood-oscillation-only model, the number of consecutive trials at which true autocorrelation coefficients exceeded the autocorrelation coefficients for randomly permuted data did not differ with respect to stimulus-congruence (2.62 ± 1.39×10^-3^ trials; T(4.32 × 10^3^) = 1.85, p = 0.06), but decreased with respect to history-congruence (2.4 ± 8.45 × 10^-4^ trials; T(4.32 × 10^3^) = −15.26, p = 3.11 × 10^-51^) relative to the full model. D. In the likelihood-oscillation-only model, the smoothed probabilities of stimulus- and history-congruence (sliding windows of ±5 trials) fluctuated as *1/f noise*, i.e., at power densities that were inversely proportional to the frequency (power ∼ 1/*f ^β^*; stimulus-congruence: *β* = −0.81 ± 1.17 × 10^-3^, T(1.92 × 10^5^) = −688.65, p < 2.2 × 10^-308^; history-congruence: *β* = −0.79 ± 1. 14 × 10^-3^, T(1.92 × 10^5^) = −698.13, p < 2.2 × 10^-308^). E. In the likelihood-oscillation-only model, the distribution of phase shift between fluctuations in simulated stimulus- and history-congruence peaked at half a cycle (*π* denoted by dotted line). In contrast to the full model, the dynamic probabilities of simulated stimulus- and history-congruence were positively correlated (*β* = 2.7 × 10^-3^ ± 7.6 × 10^-4^, T(2.02 × 10^6^) = 3.55, p = 3.8 × 10^-4^). F. In the likelihood-oscillation-only model, the average squared coherence between fluctuations in simulated stimulus- and history-congruence (black dottet line) was reduced in comparison to the full model (T(3.51×10^3^) = −4.56, p = 5.27 × 10^-6^) and amounted to 3.43 ± 1.02 × 10^-3^%. G. Similar to the full model, confidence simulated from the likelihood-oscillation-only model was enhanced for stimulus-congruent choices (*β* = 0.03 ± 1.42 × 10^-4^, T(2.1 × 10^6^) = 191.78, p < 2.2 × 10^-308^) and history-congruent choices (*β* = 9.1 × 10^-3^ ± 1.25 × 10^-4^, T(2.1 × 10^6^) = 72.51, p < 2.2 × 10^-308^). H. In the likelihood-oscillation-only model, the positive quadratic relationship between the mode of perceptual processing and confidence was markedly reduced in comparison to the full model (*β*2 = 0.34 ± 0.1, T(2.1 × 10^6^) = 3.49, p = 4.78 × 10^-4^). The horizontal and vertical dotted lines indicate minimum posterior certainty and the associated mode, respectively.

**Supplemental Figure S10.**
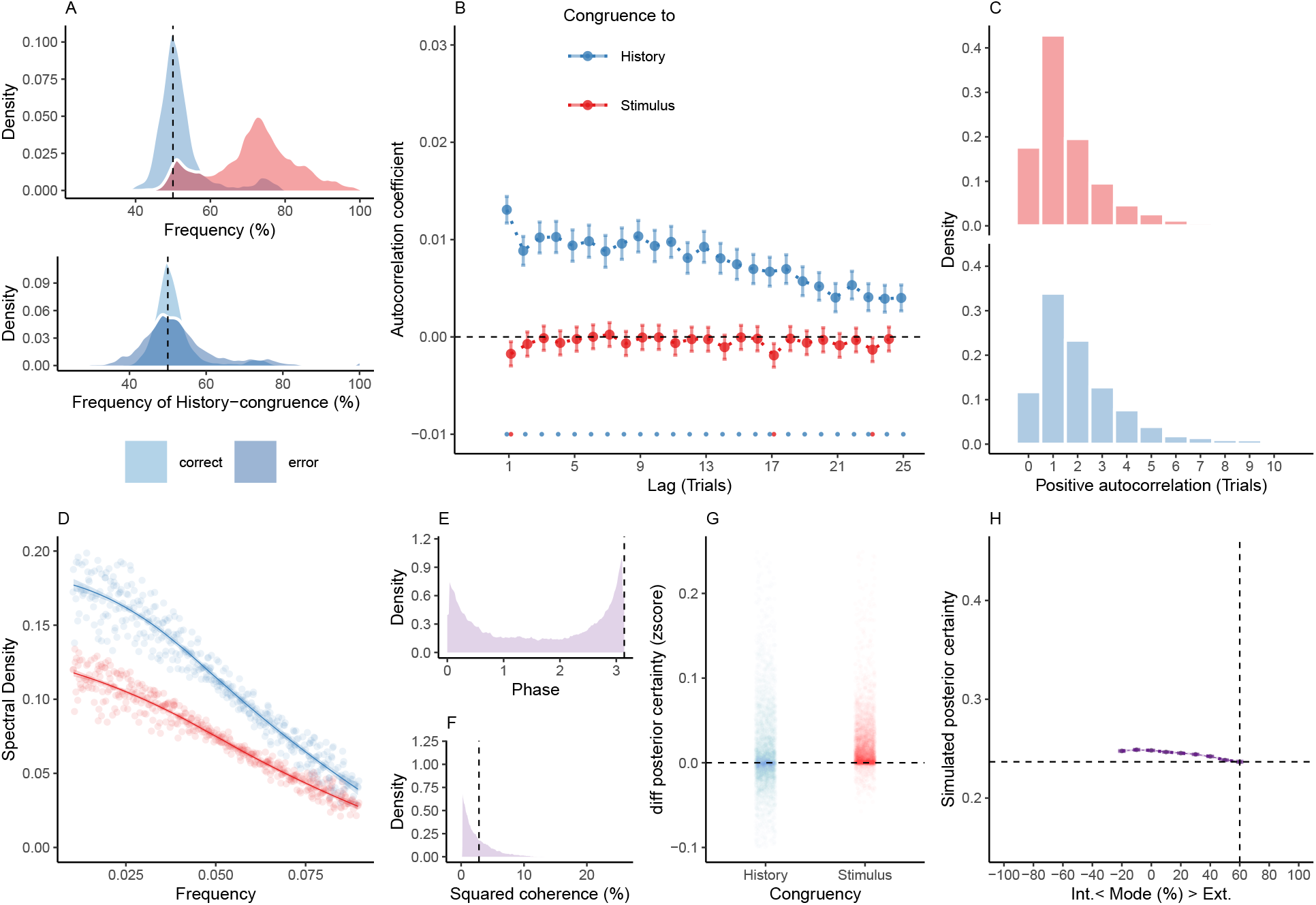
Reduced Control Model 4: Only oscillation of the prior. When simulating data for the *prior-oscillation-only model*, we removed the oscillation from the prior term by setting the amplitude *aLLR* to zero. Simulated data thus depended only on the participant-wise estimates for hazard rate *H*, amplitude *aψ*, frequency *f*, phase *p* and inverse decision temperature *ζ*. A. Similar to the full model (Figure 6), simulated perceptual choices were stimulus-congruent in 71.97% ± 0.17% of trials (in red). History-congruent amounted to 52.1% ± 0.11% of trials (in blue). As in the full model, the prior-oscillation-only showed a significant bias toward perceptual history T(4.32 × 10^3^) = 18.34, p = 1.98 × 10^-72^; upper panel). Similarly, history-congruent choices were more frequent at error trials (T(4.31 × 10^3^) = 12.35, p = 188 × 10^-34^; lower panel). B. In the prior-oscillation-only model, we did not observe any significant positive autocor­relation of stimulus-congruence, whereas the autocorrelation of history-congruence was preserved. C. In the prior-oscillation-only model, the number of consecutive trials at which true au­tocorrelation coefficients exceeded the autocorrelation coefficients for randomly permuted data did was decreased with respect to stimulus-congruence relative to the full model (1.8 ± 1.01 × 10^-3^ trials; T(4.31 × 10^3^) = −6.48, p = 1.03 × 10^-10^), but did not differ from the full model with respect to history-congruence (4.25 ± 1.84 × 10^-3^ trials; T(4.32 × 10^3^) = 0.07, p = 0. 95). D. In the prior-oscillation-only model, the smoothed probabilities of stimulus- and history-congruence (sliding windows of ±5 trials) fluctuated as *1/f noise*, i.e., at power densities that were inversely proportional to the frequency (power ∼ 1/*f ^β^*; stimulus-congruence: *β* = −0.78 ± 1.11 × 10^-3^, T(1.92 × 10^5^) = −706.62, p < 2.2 × 10^-308^; history-congruence: *β* = −0.83 ± 1.27 × 10^-3^, T(1.92 × 10^5^) = −651.6, p < 2.2 × 10^-308^). E. In the prior-oscillation-only model, the distribution of phase shift between fluctuations in simulated stimulus- and history-congruence peaked at half a cycle (*π* denoted by dotted line). Similar to the full model, the dynamic probabilities of simulated stimulus- and history-congruence were anti-correlated (*β* = −0.03 ± 8.61 × 10^-4^, T(2.12 × 10^6^) = −34.03, p = 8.17× 10^-254^). F. In the prior-oscillation-only model, the average squared coherence between fluctuations in simulated stimulus- and history-congruence (black dottet line) was reduced in comparison to the full model (T(3.54×10^3^) = −3.22, p = 1.28 × 10^-3^) and amounted to 3.52 ± 1.04 × 10^-3^%. G. Similar to the full model, confidence simulated from the prior-oscillation-only model was enhanced for stimulus-congruent choices (*β* = 0.02 ± 1.44 × 10^-4^, T(2.03 × 10^6^) = 128.53, p < 2.2 × 10^-308^) and history-congruent choices (*β* = 0.01 ± 1.26 × 10^-4^, T(2.03 × 10^6^) = 88.24, p < 2.2 × 10^-308^). H. In contrast to the full model, the prior-oscillation-only model did not yield a positive quadratic relationship between the mode of perceptual processing and confidence (*β*2 = −0.17 ± 0.1, T(2.04 × 10^6^) = −1.66, p = 0.1). The horizontal and vertical dotted lines indicate minimum posterior certainty and the associated mode, respectively.

**Supplemental Table T1.**
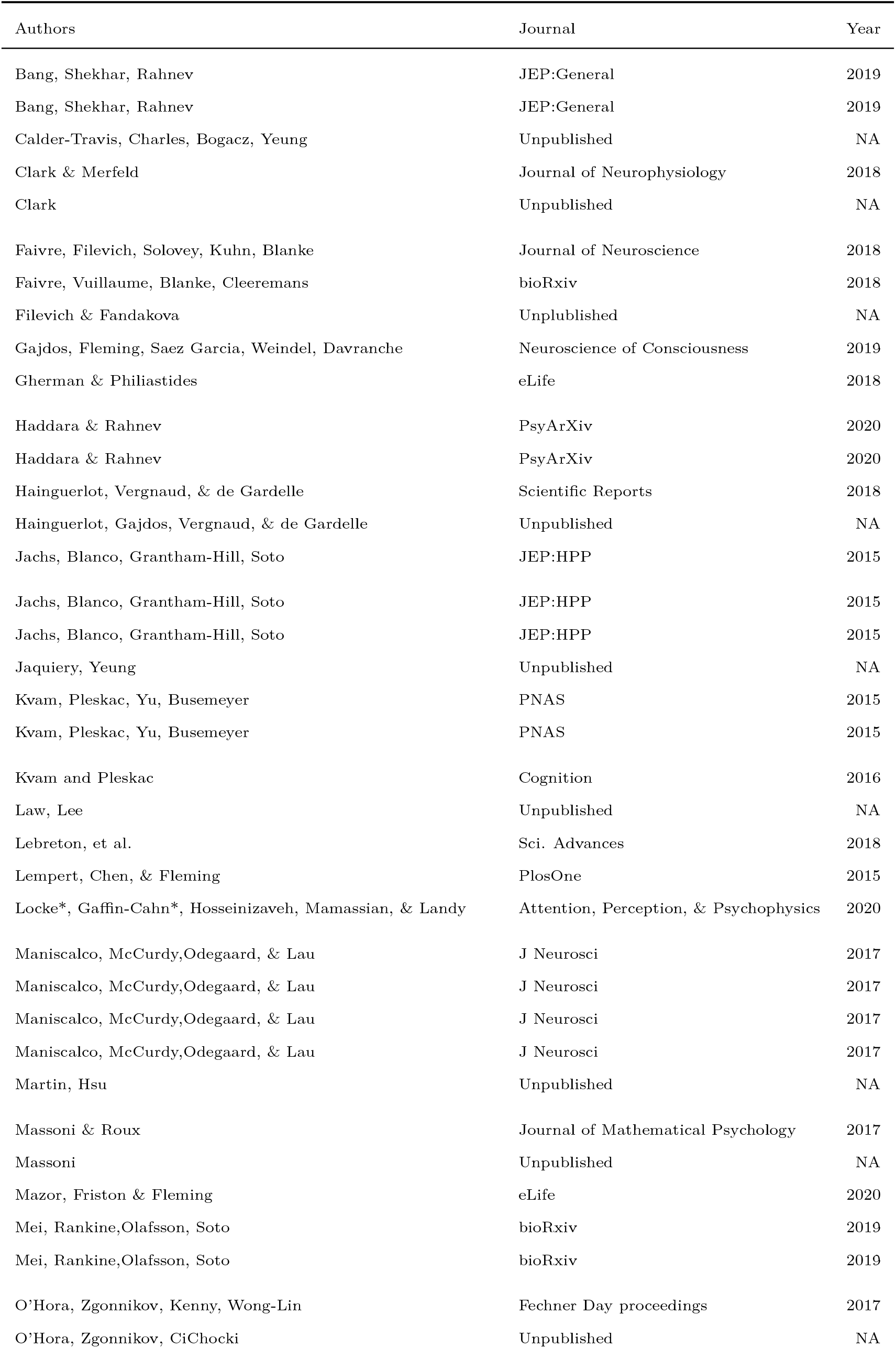
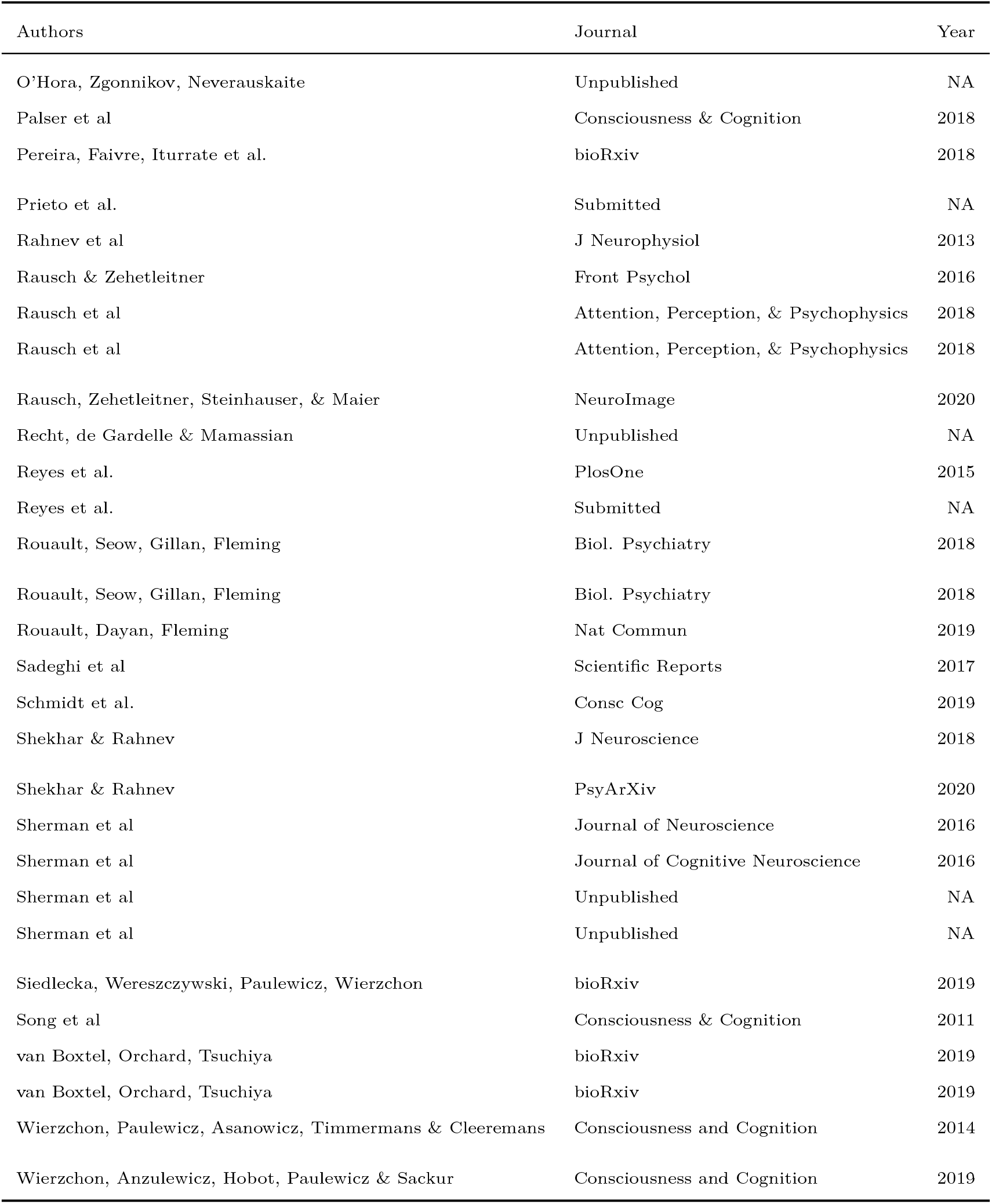

